# Dissecting infant leukemia developmental origins with a hemogenic gastruloid model

**DOI:** 10.1101/2022.10.07.511362

**Authors:** Denise Ragusa, Chun-Wai Suen, Gabriel Torregrosa-Cortés, Fabio Pastorino, Ayona Johns, Ylenia Cicirò, Liza Dijkhuis, Susanne van den Brink, Michele Cilli, Connor Byrne, Giulia-Andreea Ionescu, Joana Cerveira, Kamil R. Kranc, Victor Hernandez-Hernandez, Mirco Ponzoni, Anna Bigas, Jordi Garcia-Ojalvo, Alfonso Martinez Arias, Cristina Pina

## Abstract

Current in vitro models of developmental blood formation lack spatio-temporal accuracy and weakly replicate successive waves of hematopoiesis. Herein, we describe a mouse embryonic stem cell (SC)-derived 3D hemogenic gastruloid (haemGx) that captures multi-wave blood formation, progenitor specification from hemogenic endothelium (HE), and generates hematopoietic progenitors capable of short-term engraftment of immunodeficient mice upon maturation in an in vivo niche. We took advantage of the haemGx model to interrogate the origins of infant acute myeloid leukemia (infAML). We focused on MNX1-driven leukemia, representing the commonest genetic abnormality unique to the infant group. Enforced MNX1 expression in haemGx promotes the expansion and in vitro transformation of yolk sac-like erythroid-myeloid progenitors at the HE-to-hematopoietic transition to faithfully recapitulate patient transcriptional signatures. By combining phenotypic, functional and transcriptional profiling, including at the single-cell level, we establish the haemGx as a useful new model for the study of normal and leukemic embryonic hematopoiesis.

## INTRODUCTION

Blood formation in the embryo is a multi-stage process that develops across multiple cellular niches, reflecting a complex hierarchy of tissue interactions. It involves sequential production of distinct cell types in so-called hematopoietic waves, separated in time and space (Lacaud and Kouskoff, 2017; Costa, Kouskoff and Lacaud, 2012; Medvinsky, Rybtsov and Taoudi, 2011; Dzierzak and Bigas, 2018). The first wave of hematopoiesis produces unipotent red blood cell and macrophage precursors in the yolk sac (YS) from an angioblast precursor which can also form endothelium; it occurs at mouse embryo day (E)7-E7.5 (Fujimoto *et al.,* 2001; McGrath and Palis, 2005; Moore and Metcalf, 1970). The subsequent wave, known as definitive, produces: first, oligopotent erythro-myeloid progenitors (EMPs) in the YS (E8-E8.5); and later myelo-lymphoid progenitors (MLPs – E9.5-E10), multipotent progenitors (MPPs – E10-E11.5), and hematopoietic stem cells (HSCs – E10.5-E11.5), in the aorta-gonad-mesonephros (AGM) region of the embryo proper (Ivanovs *et al.,* 2011; McGrath *et al.,* 2015; Palis *et al.,* 1999; Medvinsky and Dzierzak, 1996; Medvinsky, Rybtsov and Taoudi, 2011). Definitive blood waves are characteristically specified from a specialised hemogenic endothelium (HE) through a process of endothelial-to-hematopoietic transition (EHT) (Zovein *et al.,* 2008; Li, W. *et al.,* 2005). HSC specification from HE is the last event in hematopoietic development and occurs in characteristic cell clusters found on the ventral wall of the dorsal aorta (Ivanovs *et al.,* 2011; Boisset *et al.,* 2010). All blood cell types enter circulation from their original locations and migrate to the fetal liver (FL) (Ciriza *et al.,* 2013; Ghiaur *et al.,* 2008). Most embryonic blood progenitors are transient and sustain blood production exclusively during embryonic development, through differentiation. In contrast, HSCs make little or no contribution to embryonic and fetal blood: instead, they proliferate (i.e. self-renew) to expand their numbers. At least a subset of HSC migrates to the bone marrow (BM) niche at the end of gestation and balances self-renewal and differentiation to maintain blood production throughout the entire post-natal life (Bowie *et al.,* 2006; Kikuchi and Kondo, 2006; Ganuza *et al.,* 2022; Yokomizo *et al.,* 2022).

Blood progenitors and HSCs can be targeted by genetic alterations that confer clonal advantage and/or drive initiation and development of leukemia. Leukemia is the most common malignancy and the most common cause of cancer-related death in the paediatric age group (Steliarova-Foucher *et al.,* 2017; Dong *et al.,* 2020). Several paediatric leukemias originate *in utero*, and a subset of these are driven by mutations exclusive to the paediatric group (Masetti *et al.,* 2015; Cazzola *et al.,* 2021). Such paediatric-exclusive mutations target blood cell types restricted to development or depend on signals that are specific to embryonic hematopoietic niches, effectively defining distinct biological entities to adult malignancies (Cazzola *et al.,* 2021; Bolouri *et al.,* 2018; Chaudhury *et al.,* 2018). Dissection of these forms of developmental leukemia is confounded by the inability to robustly identify and isolate their embryonic cells of origin, and/or to fully recapitulate the disease in post-natal cells, which leads to failure in generating relevant targeted therapies (Alexander and Mullighan, 2021). One such example is t(7;12)(q36;p13) acute myeloid leukemia (AML) driven by rearrangement and overexpression of the *MNX1* gene locus: *MNX1*-rearranged (MNX1-*r*) AML is restricted to infants under the age of 2 and carries a high risk of relapse and poor survival (Ragusa *et al.,* 2023). *MNX1* is a developmental gene required for specification of motor neurons and pancreatic acinar cells (Thaler *et al.,* 1999; Harrison *et al.,* 1999). MNX1 is not known to participate in embryonic hematopoiesis, however it has been discordantly described to be expressed in some subsets of adult hematopoietic cells (Deguchi and Kehrl, 1991; Nagel *et al.,* 2005; Wildenhain *et al.,* 2012; Taketani *et al.,* 2008). It is ectopically expressed in AML cells carrying one of a variable set of t(7;12) translocations which place the normally silent *MNX1* gene under the control of *ETV6* regulatory elements (Weichenhan *et al.,* 2023; Bousquets-Muñoz *et al.,* 2024; Ragusa *et al.,* 2023; Ballabio *et al.,* 2009). *ETV6* is required for hematopoietic cell specification from HE via VEGF signalling (Ciau-Uitz *et al.,* 2010), potentially positioning *MNX1* within a critical embryonic hemogenic regulatory network.

Faithful *in vitro* recapitulation of developmental blood cell types and their embryonic niches has the potential to dissect the cellular and molecular mechanisms presiding at initiation of developmental leukemia, reveal prognostic biomarkers, and indicate therapeutic vulnerabilities which are otherwise challenging to test in the embryo. Despite widespread use of embryonic stem (ES) cell and induced pluripotent stem (iPS) cell-based models of blood specification, a system which recapitulates the spatial and temporal complexity of embryonic blood formation within the respective time-dependent niches is still lacking (Dijkhuis *et al.,* 2023). In the last few years, gastruloid models have emerged as robust avatars of early development, self-organising processes of symmetry breaking, elongation, multi-axis formation, somitogenesis and early organogenesis, with remarkable similarity to embryo processes in space and time (van den Brink, Susanne C and van Oudenaarden, 2021; Beccari *et al.,* 2018; van den Brink, Susanne C *et al.,* 2020). Gastruloids are grown individually in a multi-well format that facilitates tracking and screening of developmental processes. Relevant to blood formation, a variant gastruloid culture system, which successful organises a heart primordium (Rossi *et al.,* 2021), was also able to recapitulate YS blood formation with erythroid and myeloid cellularity (Rossi *et al.,* 2021; Rossi *et al.,* 2022).

Herein, we describe a new protocol of gastruloid formation, i.e. hemogenic gastruloids (haemGx), that captures putative YS-like and AGM-like waves of definitive blood progenitor specification from HE. The earlier wave of putative YS-like hematopoiesis includes EMP-like cells which further mature in the gastruloid culture. HaemGx AGM-like hematopoiesis captures the emergence of intra-aortic cluster-like cells, including cells capable of short-term multi-lineage hematopoietic engraftment of the spleen and BM of immunodeficient mice. We harnessed the embryonic hematopoietic context provided by haemGx to attest its utility in modelling leukemias with developmental origins, specifically by recapitulating MNX1-*r* infant leukemia by proxy of *MNX1* overexpression. MNX1-overexpressing haemGx showed expansion of cells at the HE-to-EMP transition, which became susceptible to transformation *in vitro* and recapitulated MNX1-*r* AML patient signatures, placing *MNX1*’s leukemogenic effects at a very early stage of developmental blood formation.

## RESULTS

### HaemGxs capture time-dependent emergence of endothelium and hematopoietic cells

With the aim of recapitulating embryonic hematopoiesis with spatial and temporal accuracy, we modified existing gastruloid formation protocols to (1) promote the specification of hemogenic mesoderm derivatives, and (2) extend developmental time beyond the E8.0-equivalent of early developmental gastruloids (Van den Brink, Susanne C *et al.,* 2014) and the E9.0 timepoint of cardiogenic gastruloids (Rossi *et al.,* 2022). We established a 216h haemGx protocol (**Fig. 1A**) using a *Flk1-GFP* mouse ES cell line (Jakobsson *et al.,* 2010), which allows the tracking of early hemato-endothelial specification by expression of the *Flk1* (i.e. *Kdr*) *locus*. The haemGx protocol includes an indispensable pulse of Activin A to induce hemato-endothelial programmes via TGF-β signalling (visible by proxy of GFP-tagged *Flk1* activation), in addition of the characteristic gastruloid patterning via WNT activation by CHI99021 supplementation at 48h (**Fig. 1S1A-B**). We promoted hemato-endothelial programs through addition of VEGF and FGF2 at 72h (Sroczynska *et al.,* 2009) (**Fig. 1A**). We observed extension of a polarised *Flk1-GFP*-expressing branched endothelial network (**Fig. 1B**). Flow cytometry analysis revealed the onset of VE-cadherin^+^ C-Kit^+^ cells suggestive of EHT, with a peak at 120-144h (**Fig. 1C-D**). This was followed by a surge in CD41^+^ candidate hematopoietic cells at 144h (**Fig. 1C-D**). In our experience, observation of early polarised activation of the *Flk1 locus* at 96h (**Fig. 1B**) was predictive of CD41^+^ detection 2 days later, suggesting that TGF-β activation in early patterning is required for later hematopoietic development, putatively by modifying mesoderm specification. We incorporated a 24-hour pulse of sonic hedgehog (Shh) between 144-168h to mimic the aortic patterning that precedes cluster formation (**Fig. 1A**) (Rybtsov *et al.,* 2014). Continued exposure to VEGF after the Shh pulse resulted in specification of CD45^+^ hematopoietic cells progenitors (**Fig. 1C-E**), which were apparent after 168h. The proportion of CD45^+^ cells was enhanced by the addition of SCF, FLT3L and TPO (**Fig. 1S1C-D**), a combination of cytokines routinely used for HSC maintenance; conversely, it did not require continued addition of FGF2 after 168h (**Fig. 1S1E**), which was omitted. CD45^+^ cells, some of which co-expressed C-Kit, were present in a small number of ‘cluster’-like structures adjacent to Flk1-GFP^+^ Kit^+^ endothelium (**Fig. 1E**). The process is overall reminiscent of hematopoietic progenitor cell budding from the aortic HE, suggesting recapitulation of hemogenic tissue architecture. Time-dependent specification of C-Kit^+^, CD41^+^ and CD45^+^ cells in haemGx could be obtained with other mouse ES cell lines, including commonly used E14Tg2a (E14) cells (**Fig. 1S1F-H**), with successful specification of comparable frequencies of CD45^+^ progenitors (**Fig. 1S1I**), attesting to the reproducibility of the protocol across cell lines of different genetic backgrounds.

**Figure 1.**
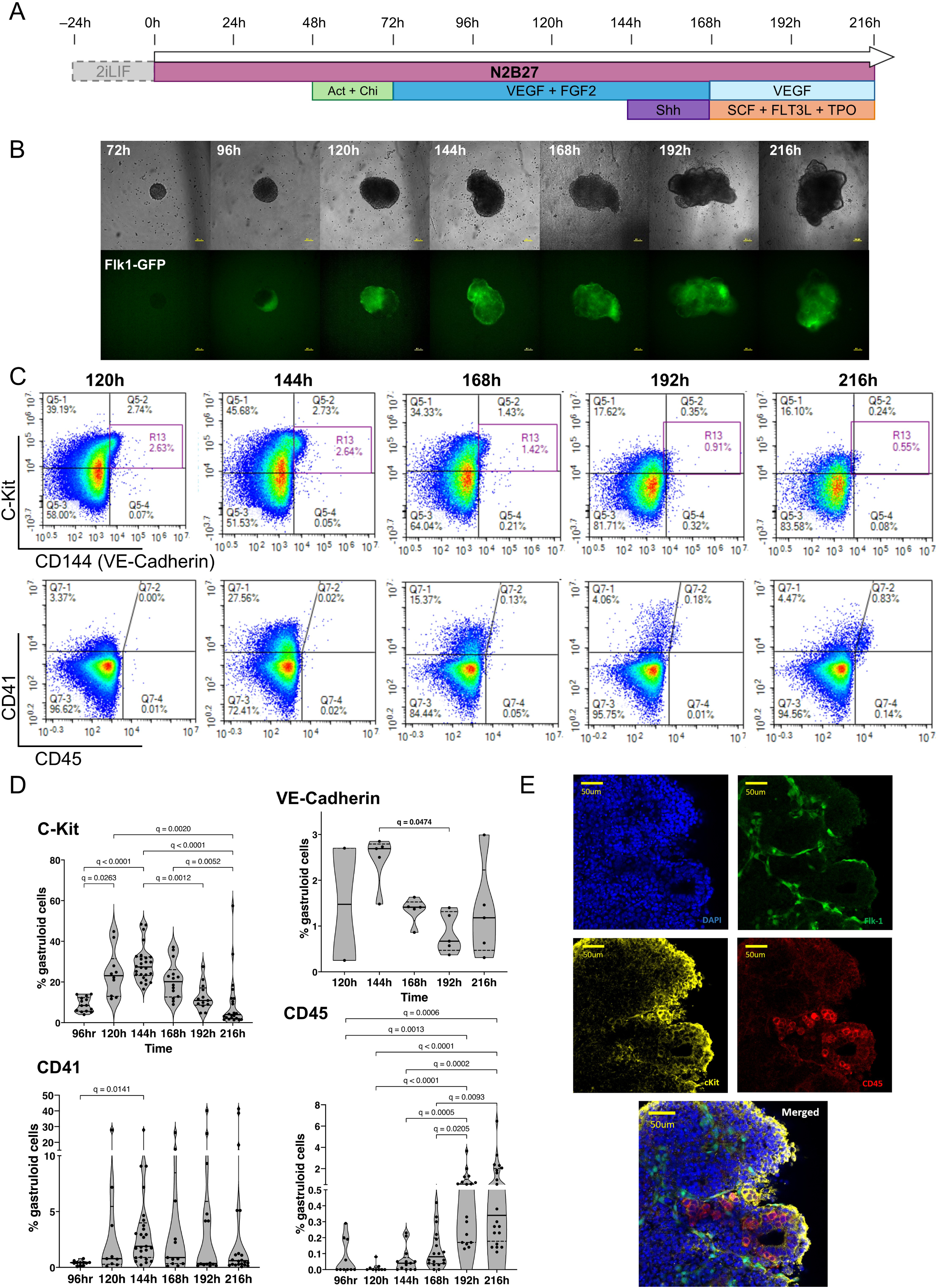
Hemogenic gastruloids (haemGx) produced from mES cells promote hemato-endothelial specification with spatio-temporally accurate ontogeny. **(A)** Timeline of mES cells assembly and culture into gastruloids over a 216h period with the addition of specified factors for the promotion of hemato-endothelial specification. The 24-hour pre-treatment in 2i+LIF was omitted when mES cultures showed no signs of differentiation. **(B)** Imaging of haemGx over time from 72h to 216h at 10x magnification, showing the assembly and growth of the 3D structures and the polarization of the Flk-1-GFP marker from 96h; scale bar: 100mm. **(C)** Flow cytometry timecourse analysis of haemGx for the presence of CD144 (VE-cadherin), C-Kit, CD41, and CD45 markers; representative plots. **(D)** Quantification of analysis in (C). Violin plots with median and interquartile range of up to 26 replicates; Kruskal-Wallis CD144 (p=0.0755), C-Kit (p<0.0001), CD41 (p<0.0342), CD45 (p<0.0001). Shown are significant Dunn’s multiple comparisons tests; adjusted q-value. **(E)** Immunostaining of whole individual haemGx at 216h showing the localized expression of Flk1-GFP (green), C-Kit (yellow), and CD45 (red) markers; nuclear staining is in DAPI (blue); scale bar: 100μm.

Overall, our extended haemGx model could capture time-dependent specification of CD41^+^ and CD45^+^ hematopoietic cells, with recapitulation of the cluster-like spatial arrangement of CD45^+^ cell emergence from HE, thus showing initial promise as a tool for the exploration of normal and perturbed developmental hematopoiesis.

### HaemGx specify cells with phenotypic and functional hematopoietic progenitor characteristics

We confirmed the induction of endothelial patterning from the Flk1-GFP population by immunofluorescence staining showing the branched vascular-like network of Flk1 co-expressing CD31 (**Fig. 2A**). Endothelial identity of Flk1-GFP^+^ populations was also detectable by flow cytometry through the presence of endothelial markers CD31 and CD34 (**Fig. 2S1A-B**). Given the co-expression of CD31 and CD34 with hematopoietic progenitors emerging from HE, we inspected CD41^+^ and CD45^+^ cells for the presence of these markers. Indeed, we found that subsets of CD41^+^ and CD45^+^ hematopoietic cells co-express CD31 and CD34 (**Fig. 2B**, **Fig. 2S1C-D**), with increased relative representation of hemato-endothelial markers within CD41^+^ cells at 144h and CD45^+^ at 216h (**Fig. 2C**), consistent with two successive waves of hemogenic emergence from HE. CD45^+^ cells from 216h haemGx also co-expressed C-Kit, Flk1-GFP and VE-cadherin, a phenotype enriched in emergent hematopoietic stem and progenitor cells (HSPC) in the AGM (**Fig. 2S1E-F**). The phenotype is detected in 1.8% of CD45^+^ cells on average (**Fig. 2S1G**), corresponding to an estimated 8 cells per haemGx. In agreement, analysis of the haemGx progenitor content at 216h using multipotential CFC assays (**Fig. 2D-E**) revealed the presence of >50 CFC/haemGx, of which an average of 7 have mixed-lineage granulocytic-monocytic-erythroid (GEM) potential, numerically matching the estimated number of HSPC. Other progenitors were predominantly granulo-monocytic (GM) and erythroid (E) progenitors (**Fig. 2D-E**). Analysis of the cellularity of dissociated haemGx at the 216h timepoint using Giemsa-Wright stained cytospins revealed differentiated cells of G, M, E and megakaryocytic lineages (**Fig. 2S2A**), putatively reflecting the progeny of EMP-like cells.

**Figure 2.**
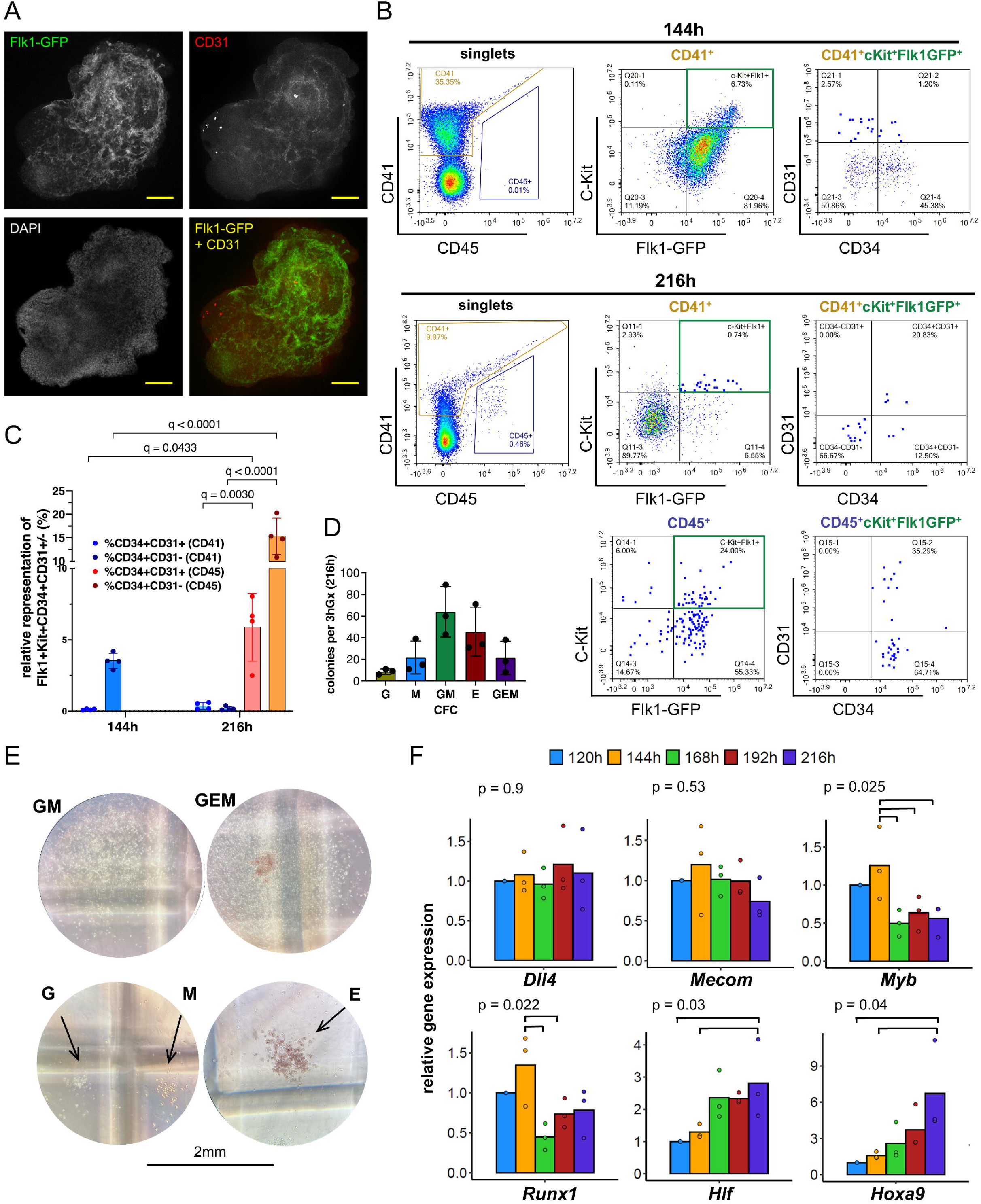
HaemGx produce hematopoietic output following the emergence of specialised endothelium. **(A)** Immunostaining of whole individual haemGx at 144h showing the establishment of a vascular network by co-expression of CD31 and Flk1-GFP; nuclear staining is in DAPI (blue); scale bar: 100μm. **(B)** Flow cytometry analysis of haemGx cells at 144h and 216h stained for CD41, CD45, CD31, and CD34. **(C)** Quantification of flow cytometry analysis in (B) of CD41+ and CD45+ fractions co-expressing CD34 and CD31. Mixed effect analysis with a Ŝidàk test for multiple comparisons. **(D)** Colony-forming cell (CFC) assay of 216h-haemGx in multipotential methylcellulose-based medium. GEM: granulocyte-erythroid-monocyte; GM: granulocyte-monocyte; M: monocyte; E: erythroid. Please note that the medium does not include TPO and does not assess the presence of megakaryocytic progenitors. CFC frequency in 3 haemGx, n=3, mean±SD. **(E)** Representative photographs of colonies quantified in C; scale bar=2mm. **(F)** Real-time quantitative (qPCR) analysis of expression of hemato-endothelial genes by timepoint. Relative gene expression fold change calculated by normalization to Ppia. Bars represent mean of 3 replicates and show individual data points; p values by ANOVA across datapoints; post-hoc statistical significance between specific variables by Tukey’s test and shown by brackets.

We observed temporal progression of expression of key markers by real-time quantitative PCR (qPCR), configuring a sustained hemato-endothelial program from 120h through 216h (*Dll4* and *Mecom*), and the generation of hematopoietic populations based on the time-dependent patterns of *Myb, Runx1*, *Hlf* and *Hoxa9* (**Fig. 2F**). We note that *Myb* peaks at 144h, coincident with the emergence of CD41^+^ cells, which we postulate to have YS-like EMP characteristics. On the other hand, *Runx1* has a bimodal pattern peaking at 144h, and later at 216h, suggesting the presence of two distinct hematopoietic waves. The later timepoint of 216h has significantly higher expression of both *Hoxa9* and *Hlf* raising the possibility that the second wave indeed corresponds to the specification of AGM-like HSPC. In order to further test the capture of YS-like and AGM-like hematopoietic waves, given the lack of markers that specifically associate with AGM hematopoiesis, we took advantage of the distinct regulatory requirements of polycomb-associated EZH2, which is necessary for EMP differentiation in the YS, but not for AGM-derived hematopoiesis or primitive erythroid progenitors (Neo *et al*., 2018; Neo *et al*., 2021). Upon EZH2 inhibition via treatment with GSK216 from 120h onwards, we observed a reduction of CD41^+^ cells selectively at 144h, with no change in CD45^+^ cells at the later timepoint of 216h (**Fig. 2S2B-C**). This suggests that the cells present at the later timepoint may not be the progeny of the former, thus supporting recapitulation of 2 independent hematopoietic waves, the latter putatively AGM. Additionally, the vulnerability of 144h CD41^+^ to EZH2 inhibition suggests that they include YS-like EMP and do not predominantly reflect EZH2-independent primitive YS hematopoiesis.

### HaemGx progenitors capture time-dependent signatures of YS-like and AGM-like hematopoiesis

To better characterize the extent and progression of developmental hematopoiesis in haemGx, we performed single-cell RNA-sequencing (scRNA-seq) time-course analysis of haemGx cell specification. We sorted cells from 2 independent haemGx cultures at 120, 144, 168, 192 and 216h, and profiled a total of 846 cells using the Smart-Seq2 protocol (Picelli *et al.,* 2014) (**Fig. 3S1A**). We analyzed unfractionated live single haemGx cells at 120-192h and enriched the dataset with sorted CD41^+^ cells at their peak time of 144h, and sorted CD45^+^ cells at 192 and 216h. At timepoints of significant enrichment in CD41^+^ (144h) and CD45^+^ cells (192h) (**Fig. 1D**), we also sorted C-Kit^+^ cells. Library preparation and sequencing generated an average of 120000 reads/cell, which were mapped to an average of 4000 genes/cell, with minimal signs of cell stress / death as measured by mitochondrial DNA fraction (**Fig. 3S1B**). Read and gene counts were similar between biological replicates, cell types and at different timepoints (**Fig. 3S1C**). The exception was 120h unfractionated haemGx cells, which despite similar sequencing depth (average read counts), were mapped to twice the number of genes (**Fig. 3S1C**), possibly reflecting multi-gene program priming at the onset of hemogenic specification. We selected highly varying genes (HVG) before principal component analysis (PCA) dimensionality reduction and retained the most relevant dimensions. In the PCA reduced space, we constructed a KNN graph and used uniform manifold approximation and projection (UMAP) to visualize the data on 2 dimensions, looking for cell communities using Leiden clustering (**Fig. 3S1D**). We identified 12 cell clusters of which 2 (clusters 0 and 5) almost exactly mapped to C-Kit^+^ cells (also positive for Sca-1), 2 (clusters 1 and 8) contained CD45^+^ sorted cells, and cluster 4 uniquely captured the CD41^+^ cells observed at 144h (**Fig. 3S1D**). Although some unfractionated cells overlapped with the hemogenic cell clusters (**Fig. 3S1D**, middle panel), most occupied different transcriptional spaces, reflecting the relative frequency of hematopoietic cells and suggesting the presence of potential hemogenic niches.

**Figure 3.**
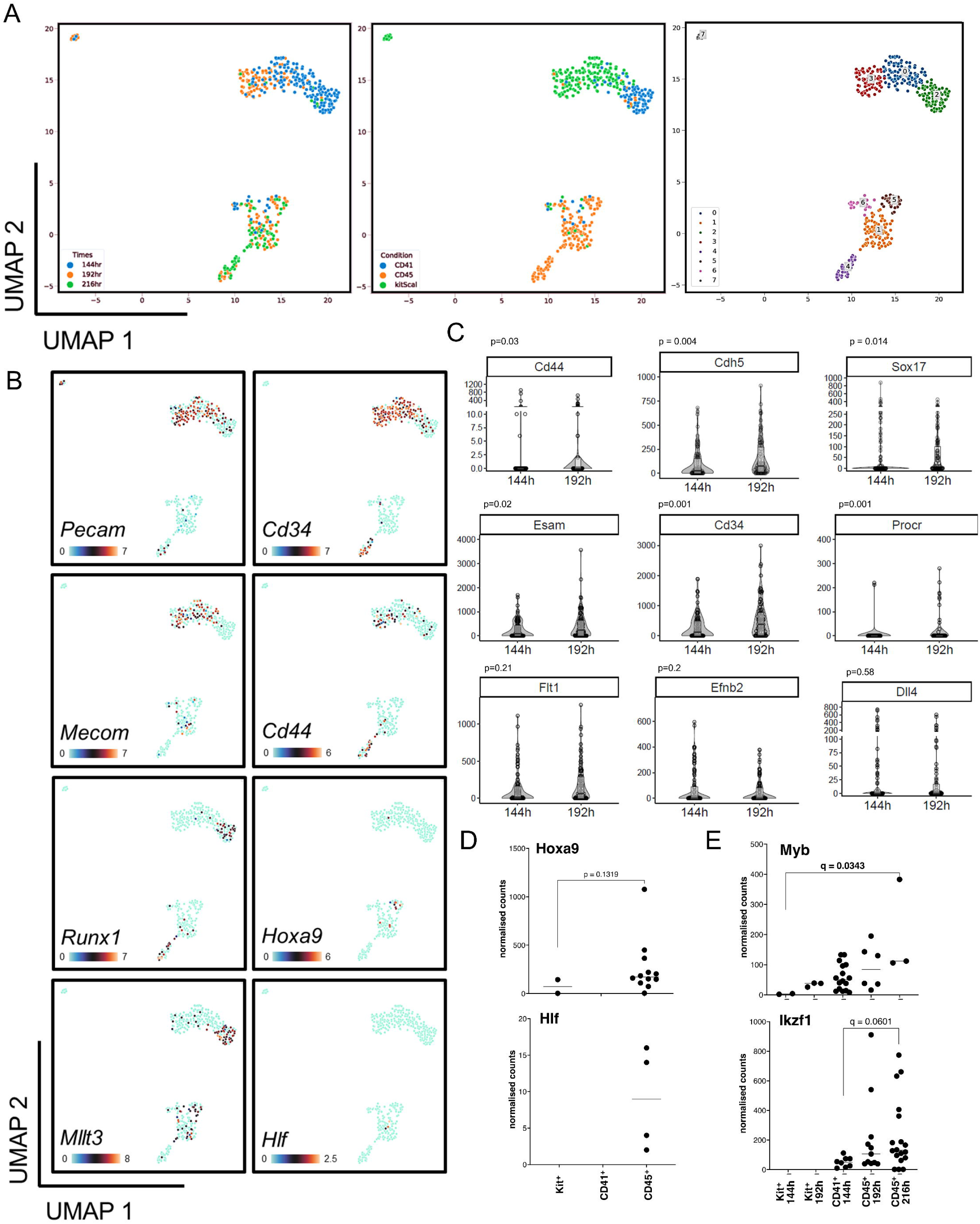
Time-resolved analysis of haemGx by scRNA-seq captures successive waves of hematopoietic specification. **(A)** UMAP clustering of subsets of sequenced cells expressing CD41, C-Kit, and CD45 at 144h, 192h and 216h, annotated by timepoint (left panel), sorting condition (middle panel), and Leiden clustering (right panel). **(B)** UMAP panels highlighting the expression of specific hemato-endothelial markers in the transcriptional spaces defined in (A). Color scale indicates expression levels in counts. **(C)** Heatmaps of differentiation of an arterial endothelial programme in haemGx from scRNA-seq data of sorted on expression of C-Kit at 144h and 192h. Color scale indicates expression in counts. **(D)** Violin plots quantifying the expression of hemato-endothelial markers in C-Kit+ fraction from clusters in (A) comparing 144h and 192h timepoints. Wilcoxon test; significant p value<0.05. **(E)** Expression of definitive hematopoietic genes in in C-Kit^+^, CD41^+^ and CD45^+^ cells at 144h and 192/216h timepoints; expression levels as normalised counts per million reads. Mann-Whitney or Kruskal-Wallis with Dunn’s multiple comparisons; significant p or q-value<0.05.

To explore tissue and lineage affiliation of the different clusters, we performed differential gene expression against all other cells using Wilcoxon ranking test, and established cluster classifier gene lists (**Supplemental File S1**). Top classifier genes for clusters 0 and 5 had endothelial affiliation (**Supplemental File S1-S2**), and the cellular composition of the clusters broadly reflected their time signature (**Fig. 3S1D**): cluster 5 comprised 144h cells and cluster 0 included most C-Kit^+^ cells at 192h. Cluster 5 occupied the vicinity of 144h CD41^+^ cells (**Fig. 3S1D**), which carried an erythroid-biased signature (**Supplemental File S2**). On the other hand, 192/216h-CD45^+^ cells in cluster 8 expressed myeloid and lymphoid genes (**Supplemental File S1-S2**). The remaining clusters largely corresponded to time-restricted unfractionated cells (**Fig. 3S1D**, bottom panel), with clusters of cells present at later timepoints matching mesenchymal stromal cell and autonomic neuron signatures (**Fig. 3S1E**), potentially configuring incipient formation of niches relevant to production of HSC (Kapeni *et al.,* 2022; Fitch *et al.,* 2012).

To specifically explore the extent to which haemGx capture putative YS and AGM-like hematopoiesis, we re-clustered cells sorted on hemato-endothelial markers CD41, C-Kit, and CD45 at the 144h, 192h and 216h timepoints (**Fig. 3A**). We mapped the expression pattern of endothelial (*Pecam* and *Cd34*), HE (*Mecom* and *Cd44*) and hematopoietic (*Runx1*, *Mllt3*, *Hoxa9*, and *Hlf*) markers across the time-dependent clusters (**Fig. 3B**). We observed a coincidence of *Pecam*, *Cd34*, and *Mecom* expression with clusters 0 and 3, which capture C-Kit^+^ cells at 144 and 192h, respectively. Some positive cells were also seen within CD41^+^ and CD45^+^-enriched clusters 2, and 1 and 4, respectively, supporting the presence of phenotypic progenitors, as suggested by flow cytometry analysis (see **Fig. 2**). CD41^+^ and CD45^+^-sorted clusters, on the other hand, show expression of *Runx1*, confirming their hematopoietic affiliation. *Cd44*, which is upregulated at the endothelial-to-hematopoietic transition (EHT) and in association with the AGM region (Oatley *et al*., 2020; Fadlullah *et al*., 2022), is detected predominantly in C-Kit^+^-enriched cluster 3 and CD45^+^-sorted clusters 1 and 4, corresponding to the later timepoints of 192 and 216h. This supports the notion that the later timepoints may capture AGM-like EHT, which is also suggested by significant increases in expression of *VE-cadherin* (*Cdh5*), *Sox17*, as well as *Cd34*, *Cd44* and *Procr* (Fadlullah *et al*., 2022; Jung *et al*., 2019; Oatley *et al*., 2020) in 192h, endothelial-enriched, C-Kit^+^ cells (**Fig. 3C-D**). Important in this context, these cells express arterial endothelial markers, including *Flt1*, *Efnb2* and *Dll4* steadily at both timepoints (**Fig. 3C and 3S2B**). *Hoxa9* expression also associates with arterial specification and definitive AGM-like hematopoiesis: albeit at low frequency, it is detected exclusively in C-Kit and CD45-sorted cells in the 192 and 216h-timepoints of haemGx differentiation (**Fig. 3B, E**). These cells also express *Mllt3* and, rarely, *Hlf*, compatible with incipient specification of an AGM-like definitive blood program (**Fig. 3B, E**). *Mllt3* is additionally expressed in CD41^+^-sorted cells of cluster 2, in line with its role in early erythroid and megakaryocytic lineage identity (Pina *et al*., 2008); *Myb* is also more frequently expressed in this population (**Fig. 3E**). In contrast, *Ikzf1*, which regulates lymphoid specification (Huang *et al*., 2019), gradually increases its expression, and indeed frequency, in 216h-CD45^+^ sorted cells. A handover from erythroid/megakaryocytic to myelo-lymphoid lineage affiliation is apparent between 144h and 216h haemGx blood cells (**Fig. 3S2C**). Of note, a small number of erythroid-affiliated cells at 216h express adult globins, suggesting a definitive origin (**Fig. 3S2C**).

Altogether, scRNA-seq analysis support the suggestion from phenotypic and *in vitro* functional data, that the haemGx model captures two waves of hematopoietic specification from HE-like cells and may configure pre-definitive and definitive embryonic blood formation.

### Late-stage haemGx cells transcriptionally resemble mouse AGM progenitors and have short-term engraftment potential

To investigate the putative YS-like pre-definitve and AGM-like definitive nature of haemGx-generated hematopoietic progenitors, we conducted projections of haemGx sc-RNAseq data onto *bona fide* embryonic populations. We considered 3 distinct scRNA-seq datasets, which (1) capture arterial and haemogenic specification in the para-splanchnopleura (pSP) and AGM region between E8.0 and E11 (Hou *et al*. 2020) (**Fig. 4S1A**); (2) uniquely capture YS, AGM and FL progenitors and the AGM EHT (Zhu *et al*., 2020) (**Fig. 4S1B**); and (3) interrogate the mouse AGM at the point of HSC emergence from the dorsal aorta HE (Thambyrajah *et al.,* 2024) (**Fig. 4S1C**). We extracted highly-varying genes from each of the 3 datasets to construct dimensionality reduction maps of the data (**Fig. 4S1A-C**), and conducted k-Nearest Neighbor (KNN) analysis to classify and regress our haemGx scRNA-seq data onto those maps considering 3 neighbors (**Fig. 4A**). HaemGx cells with correlation metric >0.7 were projected onto the respective embryonic datasets (**Supplemental File S3)**. Projections onto Hou *et al*. (2020) positioned clusters 0 and 3, corresponding to endothelium, including arterial endothelium and HE-like cells, between days E8.5 and E10, putatively spanning YS-like and AGM-like blood formation. A subset of CD45^+^-sorted cells, as well as a small numbers of CD41^+^ cells from cluster 2 projected onto E10 HE and intra-aortic cluster (IAC) cells, suggesting the presence of definitive HSPC or their precursors in haemGx (**Fig. 4A**, **Supplemental File S3**). We sought to further check the transcriptional affiliation of haemGx with intra-embryonic, definitive hematopoiesis, by considering the dataset by Zhu *et al*. (2020), which uniquely captures different hematopoietic locations in the same study (**Fig. 4A**). We matched the identity of a large proportion of CD41^+^-sorted cells to YS EMP, with some of the same cluster 2 cells mapping to erythroid progenitors, and some later CD45^+^-sorted cells resembling myeloid progenitors (**Fig. 4A**, **Supplemental File S3**). On the other hand, a subset of CD45^+^-sorted cells mapped to IAC and lymphoid-myeloid primed progenitors (LMPP) in the FL (**Fig. 4A**, **Supplemental File S3**), compatible with the presence of AGM-like definitive HSPC in haemGx. Interestingly, some CD41^+^-sorted cells also map to IAC, and indeed to HE, suggesting that this earlier population may contain HSPC precursors, in addition to YS-like progenitors. IAC and HE-like cells were also found within C-Kit^+^ cells, mostly from the 192h timepoint, while the majority of C-Kit^+^ cells captured different endothelial identities, as expected from their transcriptional profiles. Projection onto Thambyrajah *et al*. (2024) (**Fig. 4A**, **Supplemental File S3**), conflates YS-like and AGM-like progenitors onto the candidate HSC-enriched embryo AGM cluster 2, highlighting the similarity between transcriptional programs at different sites of embryonic blood emergence. CD45^+^ cells are more distally-projected onto the same cluster, as shown for E11.5 cells in the Thambyrajah *et al*. (2024) study, some of which correspond to the candidate FL-LMPP in Zhu *et al*. (2020) (**Supplemental File S3**). HaemGx cell similarities to Thambyrajah *et al*. (2024) HSC-enriched cluster 2 also include HE-like and IAC-like cells (as per Zhu *et al*. 2020 projections), mostly E9.5-like as per Hou *et al*. (2020), overall suggesting putative early specification of the AGM region within haemGx.

**Figure 4.**
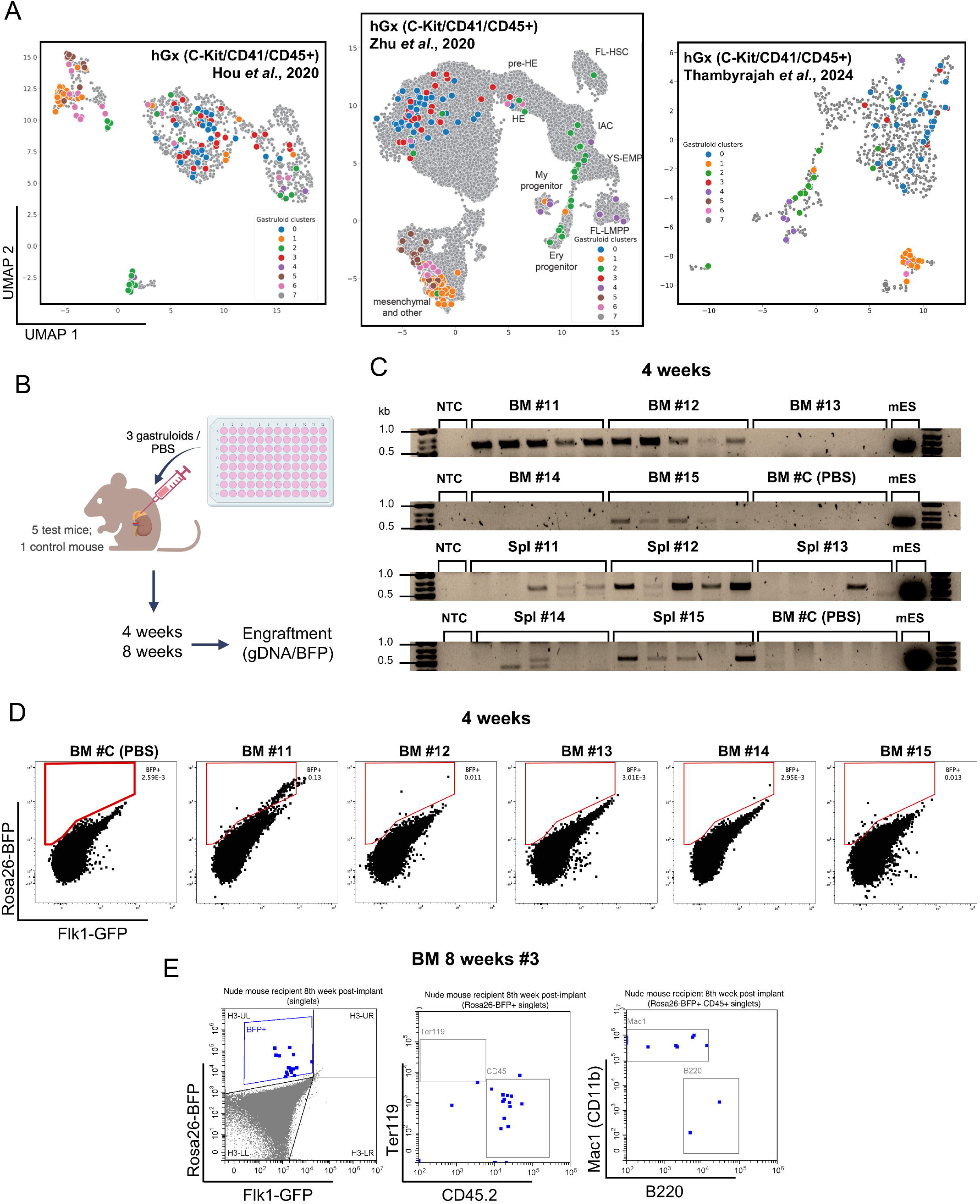
Late-stage haemGx contain hematopoietic output with transcriptional alignment to mouse embryonic populations and in vivo engraftment potential. **(A)** Projection of clustered haemGx cells (CD41, C-Kit, and CD45 at 144h, 192h and 216h; Fig. 3A) onto mouse single-cell RNA-seq datasets capturing: arterial and haemogenic specification in the para-splanchnopleura (pSP) and AGM region between E8.0 and E11 (Hou et al. 2020); YS, AGM and FL progenitors and the AGM EHT (Zhu et al., 2020); HSC emergence from the dorsal aorta HE (Thambyrajah et al., 2024). Gastruloid cell projection identified as per their cluster of origin (Fig. 3A). **(B)** Schematic representation of the experimental workflow for execution and analysis of haemGx implantation in the adrenal gland of immunodeficient mice. **(C)** PCR detection of haemGx genomic (g)DNA in the bone marrow (BM) and spleen (Spl) of immunodeficient mice 4 weeks after unilateral adrenal implantation of 3 gastruloids/gland. Analysis of 5 replicates of 100ng gDNA/recipient tissue; control animal was injected unilaterally with PBS in an adrenal gland in parallel with experimental implantation. Reaction positive control used 100ng of gDNA from *Rosa26-BFP*::*Flkj1-GFP* mES cells (mES) used to generate haemGxs. NTC: no template control. **(D)** Flow cytometry plots of engraftment detection by BFP expression in bone marrow (BM) of recipient mice 4 weeks following implantation of 216h haemGx. **(E)** Flow cytometry plots of lineage affiliation of BFP^+^ engraftment in bone marrow (BM) of recipient mice 8 weeks following implantation of 216h haemGx.

### Late-stage haemGx cells can engraft hematopoietic tissues upon maturation

The presence of putative AGM-like hematopoietic cells within haemGx led us to investigate whether haemGx-derived cells had the ability to engraft the hematopoietic system of immunodeficient animals. In early experiments, we had transplanted dissociated haemGx cells obtained at 192 and 216h into irradiated immunocompetent C57Bl/6 recipients, sub-lethally irradiated immunodeficient NSG mice, or non-conditioned NSGW41 hosts, but failed to detect signs of engraftment of peripheral blood, spleen (Spl) or bone marrow (BM) at 12 days (including absence of spleen colony formation), at 4-6 weeks (short-term engraftment), or beyond 8-10 weeks (long-term engraftment) by either flow cytometry detection of CD45.2 or by genomic DNA (gDNA) PCR analysis of the *Flk1-GFP* locus.

Failure to engraft was perhaps not surprising given that transcriptional programs position haemGx no later than E9.5-E10, which precedes HSC specification in the AGM. Additionally, specification of HSC-supporting stroma within haemGx, namely mesenchymal cells (Kapeni *et al*., 2022) and elements of the sympathetic nervous system (Fitch *et al*., 2012), is only detectable at the 192h timepoint, suggesting that further 3D maturation *in vivo* may be required. We thus tested implantation of undissociated haemGx in the adrenal gland of *Nude* mice (**Fig. 4B**), a topography more commonly used to support tumor development, but that was recently shown to be able to support HSC development, possibly with contributions from the sympathetic adrenal medullary niche (Schyrr *et al.,* 2023). In order to facilitate the analysis of engrafted animals, and given the coincidence of CD45 isoform between *Flk1-GFP* mouse ES cells and the *Nude* mice in which the experiments were performed, we engineered constitutive expression of BFP from the *Rosa26* locus in the *Flk1-GFP* mES cell line (*Rosa26-BFP::Flk1-GFP*) (**Fig. 4S2A**). We implanted 3 haemGx each unilaterally in the adrenal gland of unconditioned recipient mice (**Fig. 4B**). Control mice were injected with an equivalent volume of PBS. Animals were sacrificed at 4 or 8 weeks after implantation, with 4-5 experimental animals and 1 control collected at each timepoint. We checked Spl and BM BFP engraftment, by flow cytometry and by PCR analysis of DNA. Sensitivity of PCR analysis was determined by serial dilution of *Rosa26-BFP::Flk1-GFP* mouse ES cells in the human K562 cell line, with detection of as low as 10 pg of mouse input in a total of 100 ng gDNA (**Fig. 4S2B**), corresponding to approximately 3 cells (in 300,000). PCR analysis of BFP engraftment of the Spl and BM 4 weeks after adrenal implantation of 3 haemGx / animal detected *BFP* amplicon in both tissues in 3 of the 5 samples (#11, 12 and 15) (**Fig. 4C and 4S2C**). We sampled the gDNA of each recipient 5 times and detected the presence of the amplicon in 3-5 of the replicate samples of the positive animals; replicate sampling of the control was consistently negative (**Fig. 4C and 4S2C**). The same 3 animals showed BFP detection by flow cytometry above control, but no higher than 0.1% (**Fig. 4D**). PCR engraftment was nevertheless detectable in both erythroid Ter119^+^ and lympho-myeloid CD45^+^ fractions of the recipient animals showed PCR engraftment in both lineage fractions (**Fig. 4S2D-E**). Analysis of engraftment 8 weeks after implantation showed low-level detection (<0.1%) of BFP^+^ cells with positivity for CD11b^+^ myeloid and B220^+^ lymphoid compartments in BM of 1 recipient (**Fig. 4D and 4S2F-G**); a 4^th^ recipient had to be sacrificed early due to infection.

Overall, implantation in the adrenal niche and/or preservation of the 3D architecture enabled low-level *in vivo* hematopoietic activity, although the mechanism remains unclear. In some recipients, hematopoietic engraftment was accompanied by *in situ* growth and multi-tissue differentiation of the implanted haemGx (**Fig. 4S2H**). However, adrenal enlargement was not observed in all animals with hematopoietic engraftment (e.g. 4 weeks’ #11 had a normally sized adrenal, as did all animals analysed at 8 weeks), making it unlikely that hematopoietic activity results from teratoma formation by residual pluripotent cells. Late-stage haemGx (216h) may thus contain hematopoietic progenitors with some short-term engraftment potential but incomplete functional maturation, which are present at low frequency. Engraftment is erythro-myeloid at 4 weeks and lympho-myeloid at 8 weeks, reflecting different classes of progenitors, putatively of YS-like and AGM-like affiliation.

### HaemGx resolve the early hemogenic ontogeny targeted by infant AML gene MNX1

The ability of haemGx to reconstitute putative YS and AGM-like HE and hemopoietic progenitors positions it as an attractive model to probe cellular origins of developmental leukemia. In particular, forms of AML unique to the first 2 years of life – infant (inf)AML – have been shown to transform fetal liver (FL) but not adult hematopoietic cells (Waraky *et al.,* 2024; Mercher *et al.,* 2009; Chen, W. *et al.,* 2011; Ragusa *et al.,* 2023), compatible with targeting of a transient embryonic cell. In the case of t(7;12) AML (herein MNX1-*r*) (Ragusa *et al.,* 2023), in which the translocation results in transformation-driving ectopic expression of *MNX1* (Waraky *et al.,* 2024), analysis of patient transcriptional signatures (Ragusa *et al.,* 2022) for over-representation of cell atlas-defined affiliations, indicates enrichment in cardiomyogenic, endothelial, HSC/progenitor and mast cells (**Fig. 5A**). These signature enrichments are compatible with an early hemogenic cell derivative, capturable in the haemGx model. Other cell type-enrichments reflect MNX1 functions in pancreatic and neuronal development (Ragusa *et al.,* 2022) (**Fig. 5S1A**).

**Figure 5.**
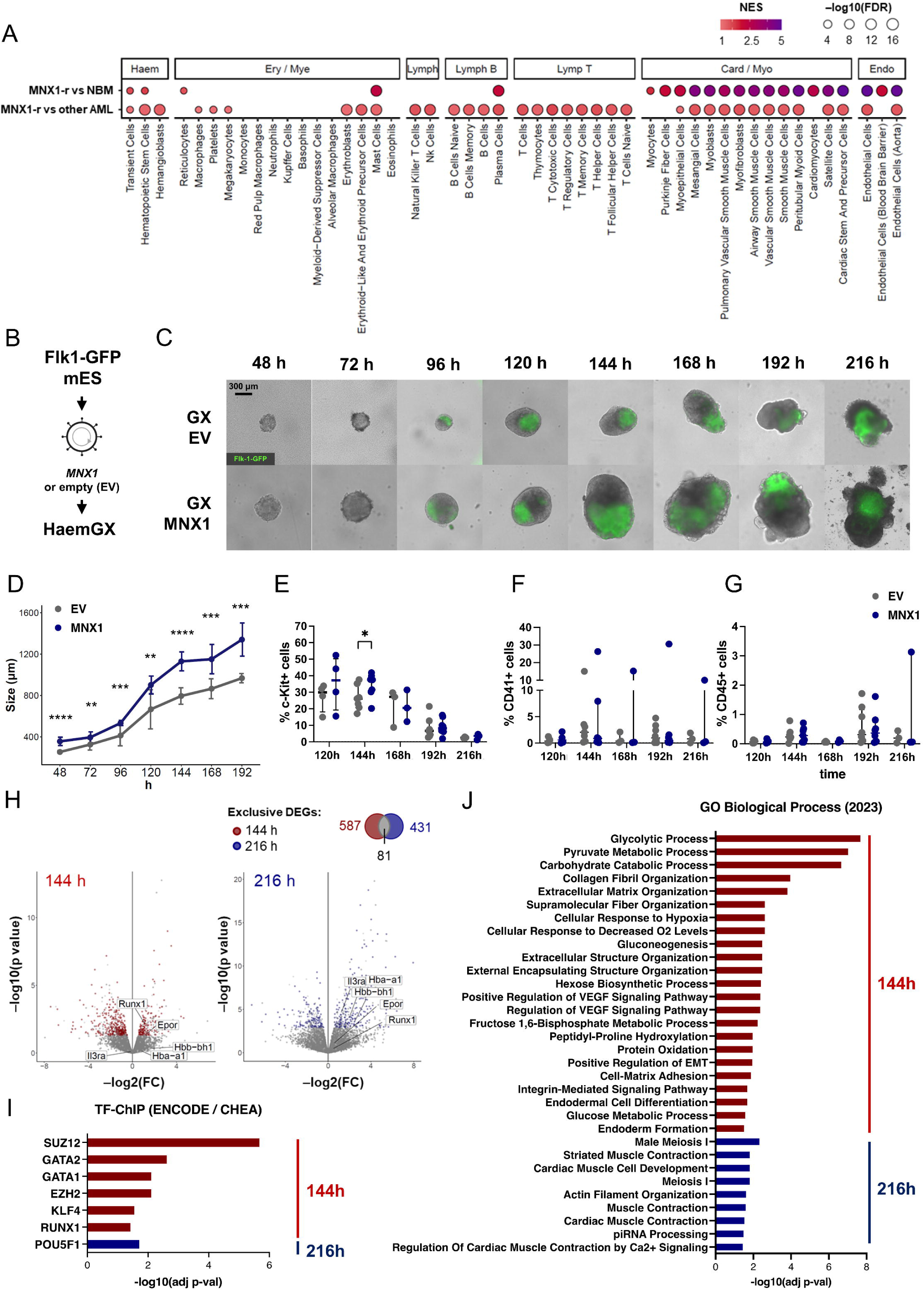
HaemGx with *MNX1* overexpression have increased cellularity and enhanced hemogenic endothelial potential. **(A)** Cell type enrichment analysis in MNX1-r AML transcriptomes, compared to normal paediatric bone marrow (BM) or other paediatric AML from the TARGET database. GSEA used representative cell type gene sets from the 2021 DB database; bubble plot shows NES scores (color gradient) and statistical significance (– log10(FDR), bubble size). **(B)** Schematic representation of generation of *MNX1*-overexpressing and empty vector (EV) mES cells by lentiviral transduction, with subsequent assembly into haemGxs. **(C)** Imaging of haemGxs with *MNX1* overexpression and EV controls at 10x magnification, showing appropriate assembly and polarization of the Flk1-GFP marker. scale bar: 300mm. **(D)** Size of haemGx at each timepoint, determined by the distance between the furthest extreme points in µm. Mean ± SD of 3 replicate experiments; 2-tailed t-test, p< 0.05 (*), 0.001 (**), 0.0001 (***), and 0.00001 (****). **(E-G)** Flow cytometry timecourse analysis of **(E)** C-Kit^+^, **(F)** CD41^+^, and **(G)** CD45^+^ cell abundance in MNX1 and EV haemGxs (120-216h). Mean ± standard deviation of 3-7 independent experiments; 2-way ANOVA and Sidak’s multiple comparison test significant at p<0.05 for C-Kit^+^ cells only (construct contribution to variance p=0.0191; 144h comparison *p=0.0190). **(H)** Volcano plots of differentially expressed genes (DEGs) by bulk RNA-seq between MNX1 and EV haemGxs at 144h and 216h. Intersection of statistically significant DEGs between 144 h and 216 h (shown in Venn diagram on top right) identified unique DEGs for each condition (labelled in red for 144 h and blue for 216 h). **(J)** Gene Ontology (GO) term enrichment for differentially upregulated genes between MNX1 and EV haemGx at 144h (red) and 216h (blue), computed in EnrichR using the GO Biological Process repository. **(I)** Transcription factor (TF) binding site enrichment on differentially upregulated genes between MNX1 and EV haemGx at 144h (red) and 216 hr (blue) using the ENCODE and ChEA Consensus TFs from ChIP database on EnrichR.

We overexpressed *MNX1* (MNX1-OE) in *Flk1-GFP* mouse ES cells by lentiviral transduction (**Fig. 5B**) and analyzed progression of haemGx generation (**Fig. 5C**). We used human *MNX1* cDNA to distinguish from the endogenous gene, but the degree of homology is nevertheless high (84%), supporting functional equivalence. We confirmed that *MNX1* overexpression in haemGx is maintained to endpoint (**Fig. 5S1B**). MNX1-OE haemGx activated polarized *Flk1-GFP* expression and elongated with similar kinetics to empty-vector (EV) control (**Fig. 5C**), albeit with formation of larger haemGx (**Fig. 5D**) denoting increased cellularity (**Fig. 5S1C**). From 192h onwards, MNX1-OE haemGx had a higher frequency of spontaneously contractile structures (**Fig. 5S1D**), compatible with the cardiogenic cell association of patients’ transcriptomes (**Fig. 5A**).

We interrogated the time-dependent hemogenic cell composition of haemGx by flow cytometry, quantifying C-Kit, CD41 and CD45 (**Fig. 5E-G**, **5S1E-F**), through which we had captured endothelial/HE-like cells, YS-EMP-like, and later putative AGM-like progenitors, respectively. We observed a small but significant expansion of the C-Kit^+^ compartment specifically at 144h (**Fig. 5E**, **5S1E**). All cell markers were relatively unchanged at later timepoints, although absolute cell numbers were higher in MNX1-OE haemGx (**Fig. 5S1C**).

To better understand the consequences of MNX1-OE on hemogenic development, we performed RNA-sequencing (RNA-seq) of haemGx at 144h and 216h, capturing putative YS and AGM-like hemopoietic waves, respectively (**Fig. 5H**). We confirmed expression of the human *MNX1* transgene in the sequenced reads (**Fig. 5S2A**); in contrast, we could not detect endogenous mouse *Mnx1* in any of the samples (**Fig. 5 S2A**), confirming that it does not normally play a role in hemogenic development. DEGs between MNX1-OE and EV (**Supplemental File S4**) configured distinct MNX1-driven programs at 144h and 216h, with >80% genes timepoint-specific (**Fig. 5H**). Both programs include up-regulation of hematopoietic-associated genes, capturing EHT and progenitor regulators (e.g. *Runx1*, and *Epor* and *Zfpm1*, respectively) at the earlier timepoint, and more differentiated effectors, e.g. *Hba-a1*, *Hbb-bh1* and *Il3ra* at 216h, compatible with selective targeting and/or expansion of YS EMP-like cells and their later differentiated progeny (**Fig. 5H**). An overview of transcriptional regulatory programs at both timepoints denotes a clearer hemogenic enrichment at 144h, with over-representation of GATA2, GATA1 and RUNX1 binding targets (**Fig. 5I**), putatively capturing an expansion of a HE-to-EMP-like transition, also represented in enriched GO categories related to VEGF signalling and epithelial-to-mesenchymal transition (EMT) (**Fig. 5J**). In contrast, no enrichment of hemogenic transcription factor (TF)-ChIP targets or related gene ontologies (GO) categories was observed in MNX1-OE haemGx at 216h (**Fig. 5I-J**), suggesting specific targeting of YS-like hemogenic programs. Instead, there was an enrichment in cardiogenic GO at 216h (**Fig. 5J**), reflecting the increase in contractile foci in MNX1-OE late haemGx.

We attempted to further deconvolve the cellular heterogeneity underlying bulk RNA-seq DEGs by interrogating the cluster signatures obtained from the single-cell analysis of haemGx development (**Fig. 5S2B-C**; refer to **Fig. 3S1D**). In support of an association with hemato-endothelial specification, MNX1-OE transcriptomes showed enrichment of haemGx HE cluster 5 and 0 signatures; the YS-EMP-like cluster 4 was also enriched (**Fig. 5S2B**). MNX1-OE DEGs at both timepoints also enriched for the relatively less differentiated cluster 10, which predominantly captures unfractionated haemGx cells at 120h, supporting an early ontogenic affiliation. Significantly, clusters 5 and 0 HE signatures were also enriched in *MNX1*-r infAML patient signatures (Ragusa *et al*., 2022) (**Fig. 5S2B-C**), putatively aligning MNX1-OE effects with HE and the hemogenic transition. In contrast, MNX1-OE DEGs specific to 216h showed enrichment in myeloid progenitor / MLP-enriched cluster 8 which captures 192/216h haemGx cells, but diverges from *MNX1*-r infAML signatures (**Fig. 5S2B-C**), instead approximating the distinct myelo-monocytic and/or mixed-lineage affiliation of *KMT2A*-rearranged AML, which is also common in older pediatric and adult patients.

### Serial replating of MNX1-OE haemGx cells identifies a candidate C-Kit^+^ leukemia-propagating cell more susceptible to transformation in the YS period

We further investigated the potential of MNX1-OE haemGx to reflect *MNX1*-r AML biology by performing serial replating of CFC assays, a classical *in vitro* assay of leukemia transformation. We compared 144h and 216h haemGx to match functional and transcriptome data, and plated unfractionated haemGx, given the transient nature of C-Kit^+^ enrichment. Both 144h (**Fig. 6A-B**) and 216h (**Fig. 6C-D**) MNX1-OE haemGx replated for at least 5 platings, significantly different to control (EV) (**Fig. 6B,D**). EV-haemGx replating could not be sustained beyond plate 3 in most experiments started from 216h (**Fig. 6C**); 144h EV-haemGx exhibited some low-level replating until plate 5 (**Fig. 6A**), but the colonies were formed by few scattered cells clearly distinct from the well-demarcated colonies obtained from MNX1-OE haemGx. We analyzed early-stage (plate 1) and late-stage (plate 5) colonies by flow cytometry to identify the nature of the replating cells. Cells selected through replating were C-Kit^+^ (**Fig. 6E**), matching the cell phenotype transiently enriched at 144h. MNX1-OE C-Kit^+^ cells co-expressed CD31 (**Fig. 6S1A-B**), underpinning the endothelial signatures of *MNX1-r* leukemia. Early replating of MNX1-OE haemGx also enriched for erythroid-affiliated Ter119^+^ cells, particularly when initiated from the 144h timepoint (**Fig. 6S1C**), but this was not sustained through replating. Myelo-monocytic-affiliated markers, including CD11b or CD45, were not present (**Fig. 6E**, **6S1C**).

**Figure 6.**
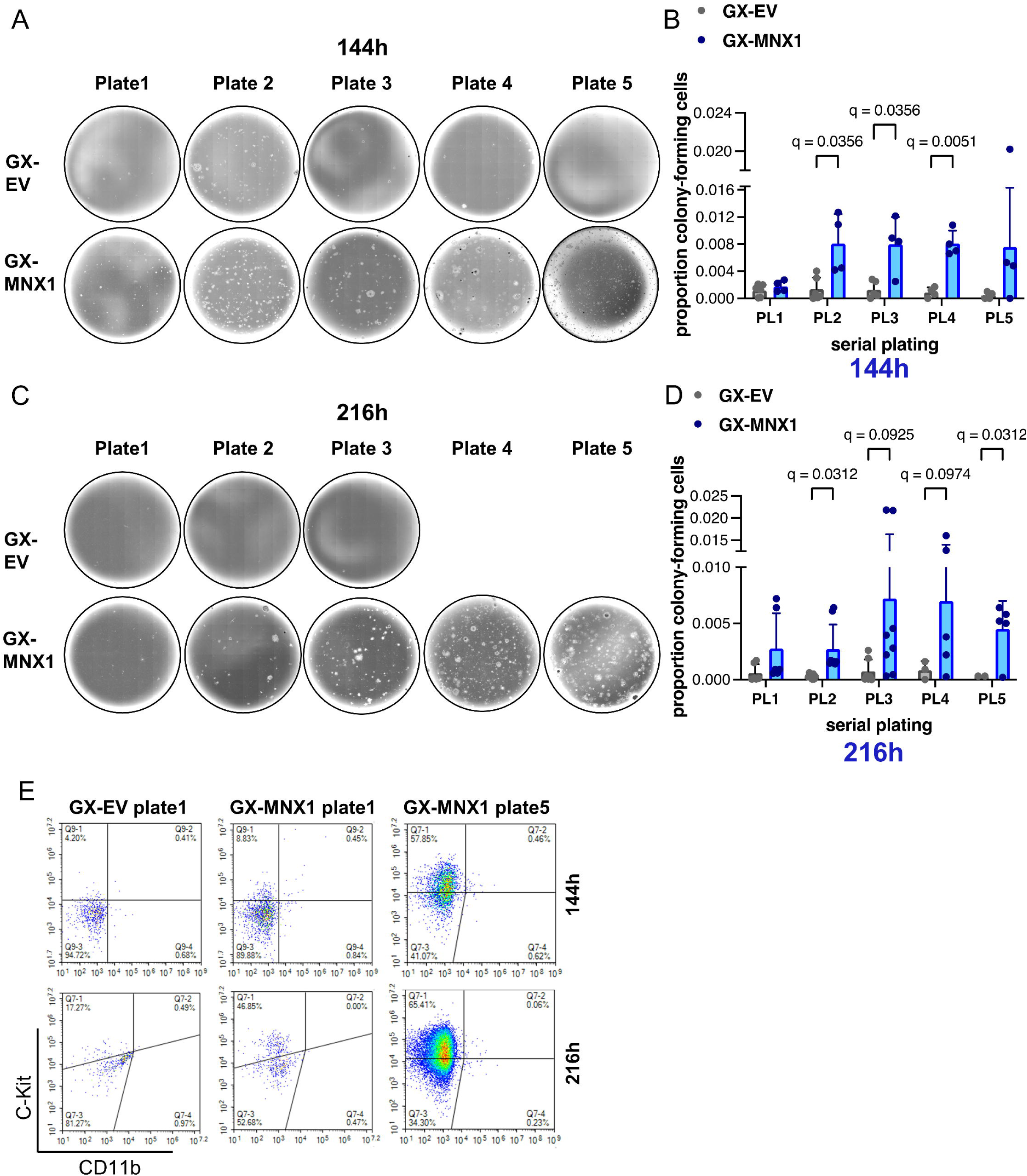
MNX1 selects C-Kit^+^ clonogenic cells and selectively transforms end-stage haemGx. **(A)** Representative photographs of serial replating of colony-forming cell assays initiated from EV or MNX1 cells obtained at 144h of the haemGx protocol. **(B)** Quantification of colony-replating efficiency of EV and MNX1 144h-haemGx (GX) cells. Mean+SD of n>3 replicates; Kruskal-Wallis with Dunn’s multiple comparison testing at significant q<0.05. **(C)** Representative photographs of serial replating of colony-forming cell assays initiated from EV or MNX1 cells obtained at 216h of the haemGx protocol. **(D)** Quantification of colony-replating efficiency of EV and MNX1 216h-haemGx cells. Mean+SD of n=5-8 replicates; Kruskal-Wallis with Dunn’s multiple comparison testing at significant q<0.05. **(E)** Flow cytometry analysis of hemogenic progenitor C-Kit^+^ cells and myeloid-affiliated CD11b^+^ cells at first round of plating of 144h and 216h-haemGx initiated from EV and MNX1 cells; representative plots.

We performed RNA-seq analysis of MNX1-OE haemGx CFC cells obtained through re-plating and integrated the data with the 144h and 216h haemGx timepoints to identify genes enriched through the process of transformation. Hs*MNX1* reads were increased in MNX1-OE CFC samples, supporting selective expansion of transduced cells (**Fig. 7A**). We performed hierarchical clustering of all DEGs in pairwise comparisons (**Fig. 7B**), and used Gene Set Enrichment Analysis (GSEA) of *MNX1*-r patient signatures up-regulated in comparison to other AML in the infant (0-2) age group, to interrogate the different clusters. Cluster k14 had significant enrichment of MNX1-*r* signatures against all other AML subtypes (**Fig. 7C**), with considerable overlap between leading-edge (i.e. significantly-enriched) genes (**Fig. 7S1A**). Importantly, k14 captures the subgroup of DEGs up-regulated upon CFC replating which are also up-regulated in MNX1-OE haemGx at 144h (**Fig. 7B**), matching the time-dependent enrichment of the putative MNX1-OE propagating population. Interrogation of cell-type-specific signatures within the k14 MNX1-*r* leading edge genes against the single-cell PanglaoDB atlas captured similarities to hemogenic and non-hemogenic cell types, including interneuron, and other neural signatures (**Fig. 7D**), which reflect MNX1 biological functions (Harrison *et al.,* 1999; Thaler *et al.,* 1999; Ragusa *et al.,* 2022). The most highly significant signatures capture early developmental hemogenic components, such as ‘endothelial cells’ and ‘erythroid cell precursors’, reflecting the 144h, putative YS affiliation of MNX1-OE haemGx. The ‘mast cells’ affiliation of *MNX1*-r AML signatures within the haemGx k14 cluster is also likely to reflect a YS EMP-like affiliation (Chia *et al*., 2023), and is accordingly aligned with MNX1-OE haemGx isolated at 144h (**Fig. 7S1B**). Giemsa-Wright-stained cytospins of serially-replated CFC assays initiated with 144h MNX1-OE cells showed a shift from blast-like at plate 3 to precursor cell morphology with some differentiated mast cells at plate 5 (**Fig. 7E, S1C**). We also observed mast cells in 216h-initiated colonies (**Fig.7E**), but these were differentiated at an earlier plate, suggesting that MNX1-OE-responsive C-Kit^+^ progenitors experience maturation between 144h and 216h.

**Figure 7.**
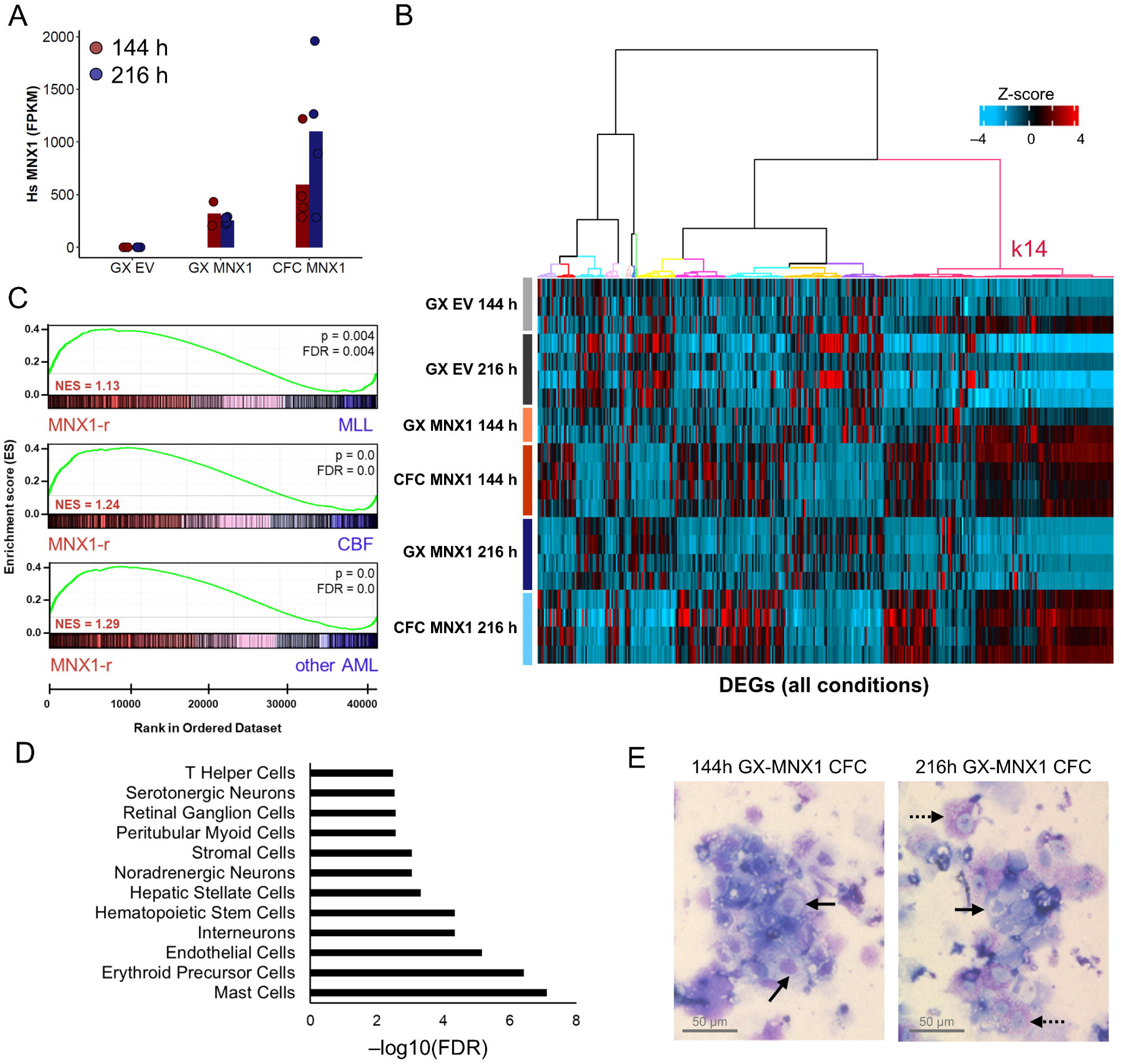
HaemGx with *MNX1* overexpression recapitulate MNX1-Acute Myeloid Leukemia patient signatures. **(A)** Quantification of human *MNX1* transcripts in FPKM units from RNA-seq of GX-EV, GX-MNX1, and CFC at 144 hr and 216 hr. Bars represent mean of replicates. **(B)** Heatmap comparing the expression of all differentially expressed genes (DEGs) between all conditions (GX-MNX1 144 hr vs GX-EV 144 hr; GX-MNX1 216 hr vs GX-EV 216 hr; CFC MNX1 144 hr vs GX MNX1 144 hr; CFC MNX1 216 hr vs GX MNX1 216 hr) as Z score. Hierarchical clustering by Ward D method on Euclidean distances identifies 14 clusters (k). **(C)** GSEA plots for gene set extracted from k14 from **(B)** against MNX1-r patients RNA-seq counts vs MLL, core-binding factors (CBF), or other pediatric AML. **(D)** Cell type analysis of the intersect of leading edge genes LEGs (n=476) from Fig. S5D in MNX1-r patients (k14 LEGs) **(B)** using the Panglao DB 2021 database. **(E)** Representative Giemsa-Wright stained dissociated CFC replating (4^th^ plating) MNX1 haemGx cells from 144h and 216h. Solid black arrows, mast cell precursors; dashed black arrows, mast cells.

Taken together, the data positioned the initiating cellular targets of MNX1-*r* AML at the YS endothelial-to-EMP transition. More broadly, the analysis supports the potential of haemGx to dissect the ontogeny of specific subtypes of hematological cancers initiated *in utero*.

## DISCUSSION

In this study, we developed and characterized a hemogenic gastruloid (haemGx) model from mouse ES cells, which recapitulates YS-like and AGM-like waves of definitive blood progenitor specification, and has incipient ability to engraft adult hematopoietic tissues short-term upon *in vivo* maturation. By exploring the haemGx model through enforced expression of the *MNX1* oncogene, which is a hallmark of infant AML harbouring the t(7;12) translocation, we identified the YS endothelial-to-hematopoietic transition as the transformation target of *MNX1* gene rearrangement, shedding light on to its exclusive association with the infant age group.

Our data positions the haemGx model as an advanced platform to probe physiological and pathological development of the hematopoietic system, that benefits from the time-dependent and topographic organization associated with gastruloid models. We constructed the haemGx model by introducing a pulse of Activin A and extending the protocol under hematopoiesis-promoting cytokines, to reach the equivalent of mouse embryonic day 10 (E10) and achieve sequential acquisition of YS-like and AGM-like blood production from HE, benchmarked by surface markers CD41 and CD45, respectively. In agreement with recent observations from Dias *et al*. (2025), which highlight a critical balance between WNT-driven posterior and Nodal / TGFβ anterior signals in patterning the midbody axis and endo-mesodermal derivatives, we observed that the combination of Activin A and CHIR99021 was critical for robust *Flk1*/*Kdr* activation and the efficient initiation of CD41 and CD45-marked hematopoietic specification. This is in contrast with the hematopoietic output of cardiogenic gastruloids reported by Rossi *et al*. (2022), which did not require Activin A. Although our strict dependence on Activin A to achieve initial activation of *Flk1* programs and specification of a vascular network and hematopoietic programs precluded a side-by-side comparison with the Rossi protocol (Rossi *et al.,* 2022), we note their apparent lower efficiency in blood formation, at least at the comparable timepoints of 144 and 168h, and their bias towards erythroid formation, reflecting a putative YS, and potentially primitive, nature. This bias may also be captured in the reported spleen colony formation; however, given the limited characterisation of the colonies in the manuscript, which does not appraise origin, morphology or cellularity, the nature of the spleen engraftment remains speculative. Importantly, standard gastruloid protocols with only CHIR99021 pulse and no additional cytokines (Beccari *et al.,* 2018) can capture a transcriptional erythrogenic wave.

Despite low frequency engraftment of adult hematopoietic tissues by haemGx cells, we demonstrated donor origin of the engrafted cells by flow cytometry and PCR detection of a BFP transgene up to 8 weeks. Engraftment progressed from erythroid-biased at 4 weeks to myelo-lymphoid at 8 weeks, as observed in progenitor engraftment from adult cells (Chun *et al*., 2024). Critically, successful engraftment required implantation of undissociated haemGx for further maturation in a supportive microenvironment, in this case provided by the adrenal gland (Schyrr *et al.,* 2023). In this study, we did not dissect the relative contribution of the continued maturation in 3D *vs* the influence of the niche for implantation, as we opted for performing the transplantation as to maximise both. Future experiments will be necessary to understand possible extrinsic niche contributions, including the sympathetic nervous system we sought to include (Kapeni *et al.,* 2022; Fitch *et al.,* 2012), and how they can be incorporated in the haemGx culture by extension and/or further modification of the protocol. Although we observed continued growth and maturation of the haemGx *in situ* after implantation, this was not the case in all animals with engraftment. On the one hand, this dissociation excludes the likelihood of the engraftment resulting from teratoma formation from the gastruloid itself. On the other hand, it raises the possibility that the inefficiency of the engraftment is the consequence of a low frequency of cells within each haemGx with HSPC properties, which we can seek to resolve by upscaling of the haemGx culture or further expansion of dissociated HSPC (Ng *et al*., 2024).

Although the system is shy of achieving HSC maturation *in situ*, it can mature, to produce low-level engrafting multilineage progenitors, and potentially HSCs. This suggests that access to the relevant hematopoietic molecular programs is preserved. Incipient specification of non-hematopoietic tissues shown to support HSC emergence, including PDGFRA^+^ mesenchymal (Chandrakanthan *et al.,* 2022) and neuro-mesenchymal (Miladinovic *et al.,* 2024) cells, and neural-crest derivatives (Kapeni *et al.,* 2022; Fitch *et al.,* 2012), occurs in sync with development of AGM-like hematopoiesis, to indicate the presence of cues that with further maturation may facilitate HSPC specification at the appropriate developmental time. Interestingly, through extending the haemGx culture, and in addition to AGM-like hematopoiesis, we also captured the potential to form renal tubular structures upon *in vivo* maturation, suggesting induction of lateral plate and intermediate mesoderm from which the AGM region derives, and putative reconstitution of the intra-embryonic hematopoietic niche.

Establishment of a tractable *in vitro* model of hematopoietic development which faithfully recapitulates time-dependent emergence of hematopoietic progenitors in their physiological niche would propel translational applications by enabling high-throughput studies of HSC emergence. Most ESC-based models of blood specification do not achieve concomitant recapitulation of the surrounding microenvironment. Protocols typically remove ESC-derived hemogenic cells from an early embryonic-like niche to direct differentiation through genetic manipulation (Sugimura *et al.,* 2017) and/or empirically-determined growth factor addition (Lis *et al.,* 2017; Vo and Daley, 2015; Fowler *et al.,* 2024; Nafria *et al.,* 2020; Sroczynska *et al.,* 2009; Piau *et al.,* 2023), with minimal probing of intermediate steps. When hematopoietic waves have been specifically inspected, emergence of YS and AGM-like stages were conflated in time (Pearson *et al.,* 2015; Murry and Keller, 2008), suggesting a bias towards intrinsic hematopoietic cell potential, at the expense of cell fate modulation by time-coherent extrinsic physiological cues, affecting the nature of cellular output. The present haemGx system, in contrast, successfully deconvolutes YS-like and AGM-like stages of blood production, including capturing differences in the HE generation, suggesting a level of organisation amenable to further development into an integrated hemogenic niche. Indeed, projections to scRNA-seq datasets of *in vivo* murine hematopoiesis place our endothelial and hematopoietic outputs at coherent temporal windows.

The haemGx protocol could be further developed to include more supportive structures of tissue organisation, as non-hematopoietic gastruloid models have benefitted from mechanical, or chemo-mechanical, support of artificial ECM embedding (Veenvliet *et al.,* 2020; van den Brink, Susanne C *et al.,* 2020; Dijkhuis *et al.,* 2023), to match temporal coherence and tissue polarisation, with detailed tissue architecture illustrated by somite formation. Prior to HSC emergence in the AGM, critical signals such as SCF and Shh, which we have delivered to the cultures, are secreted by different cells on opposite dorsal-ventral locations (Souilhol *et al.,* 2016), indicating the potential relevance of tissue organisation to improve and potentially extend hematopoietic formation. Moreover, HSPC formation in the AGM is facilitated by circulatory shear stress (North *et al.,* 2009; Diaz *et al.,* 2015; Lundin *et al.,* 2020), suggesting that introduction of flow may improve hemato-endothelial organization and boost blood production.

We attested the utility of haemGx’s to interrogate the biology of leukemias with developmental origins, thanks to a nearly full recapitulation of embryonic hemogenic specification, including critical early stages of HE production and EHT to EMP and MLP/MPPs. We modelled *MNX1*-r AML, a poorly characterized form of leukemia exclusive to the infant age group (Ragusa *et al.,* 2023). Overexpression of *MNX1* in the haemGx model propagated cells bearing transcriptional resemblance to patient *MNX1*-r AML, overall positioning the haemGx as an attractive tractable model in which to study developmental leukemias. We showed that MNX1-OE transiently expanded a C-Kit^+^ cell with hemato-endothelial characteristics at the YS-like stage, and could selectively sustain the phenotype through serial replating of CFC assays. Replating CFCs could be initiated beyond the YS-like stage, albeit at the relative expense of an early differentiation state and of the similarity to patient transcriptome, putatively defining a window of susceptibility to MNX1-driven transformation.

Importantly, the haemGx model was able to resolve the cellularity and transcriptional features inferred from patient transcriptomes, with the advantage of a higher resolution in dissecting the contribution of cell-of-origin in the appropriate developmental context. A recent paper using MNX1-OE has shown that MNX1 can initiate a transplantable leukemia in immunodeficient mice by targeting FL, but not BM, cells (Waraky *et al.,* 2024). This suggests that ectopic expression of *MNX1* transforms an embryonic cell, rather than depends on an embryonic niche. It also suggests that MNX1-OE can cause leukemia as a single genetic hit, although it cannot be excluded that other mutations are required for, and may indeed have contributed to, full transformation *in vivo*, as observed in a recent report confirming the *in utero* origin of the leukemia (Bousquets-Muñoz *et al.,* 2024). However, the mouse model does not address the nature of the embryonic cell targeted by MNX1-OE. Like the MNX1-OE haemGx cells selected through serial replating, MNX1-OE mouse leukemia are also C-Kit^+^ (Waraky *et al.,* 2024). Interestingly, clusters of genes up-regulated in the mouse MNX1-OE leukemia (**Fig. 7S1D** – k1, 2, 5, 6, and 9) do not robustly separate *MNX1*-r from other infAML by GSEA analysis (**Fig. 7S1E**), and have diverged more from the patient data in their cell-type signature analysis than the original MNX1-OE cells in the FL (**Fig. 7S1F**), putatively selecting a more generic program of leukemia transformation. With the resolution provided by the haemGx model, we propose that the divergence from *MNX1*-r infAML signatures increases from MNX1-OE haemGx 144h, to 216h to FL, positioning the origin of the AML at the YS stage, likely at the HE-to-EMP transition. This cell may persist in the haemGx culture system to the 216h timepoint, and migrate to the FL in the embryo, eventually extinguishing through differentiation, as suggested by the MNX1-OE haemGx CFC assays. Persistence of the original MNX1-OE target cell will determine the sensitivity to MNX1-OE, and can be addressed in the haemGx system through lineage tracing. Whether additional mutations, namely a reported association between *MNX1*-r and tri(19) (Espersen *et al.,* 2018; Ragusa *et al.,* 2023), contribute to prolonging the undifferentiated state of the candidate YS target cell with increased probability of transformation, or otherwise expand the targeted cell in a more classical second-hit fashion, can potentially also be screened in the haemGx system, in combination with transplantation experiments.

Over the last decade, the effort to achieve robust hematopoietic production from PSC, mouse and human, has been centred on the generation of HSC, a critical translational need (Chang *et al.,* 2023; Dijkhuis *et al.,* 2023; Fidanza and Forrester, 2021). There are good indications that this is achievable, both with (Vo and Daley, 2015; Dang *et al.,* 2002) and without (Piau *et al.,* 2023) genetic manipulation. However, success remains episodic, mainly because mechanistic understanding of processes underpinning blood generation during development is limiting, exacerbated by limiting access to human embryonic and fetal material. On-going optimization of gastruloid models and the success at integration of organoid cultures in microfluidic systems (Saorin, Caligiuri and Rizzolio, 2023; Quintard *et al.,* 2024) will allow us to further optimize haemGx culture. In conclusion, the haemGx represents a faithful *in vitro* reference to replace and complement embryo studies, holding promise to understand the biology of healthy and malignant developmental hematopoiesis and advance blood production for clinical use.

## EXPERIMENTAL PROCEDURES

### MATERIALS

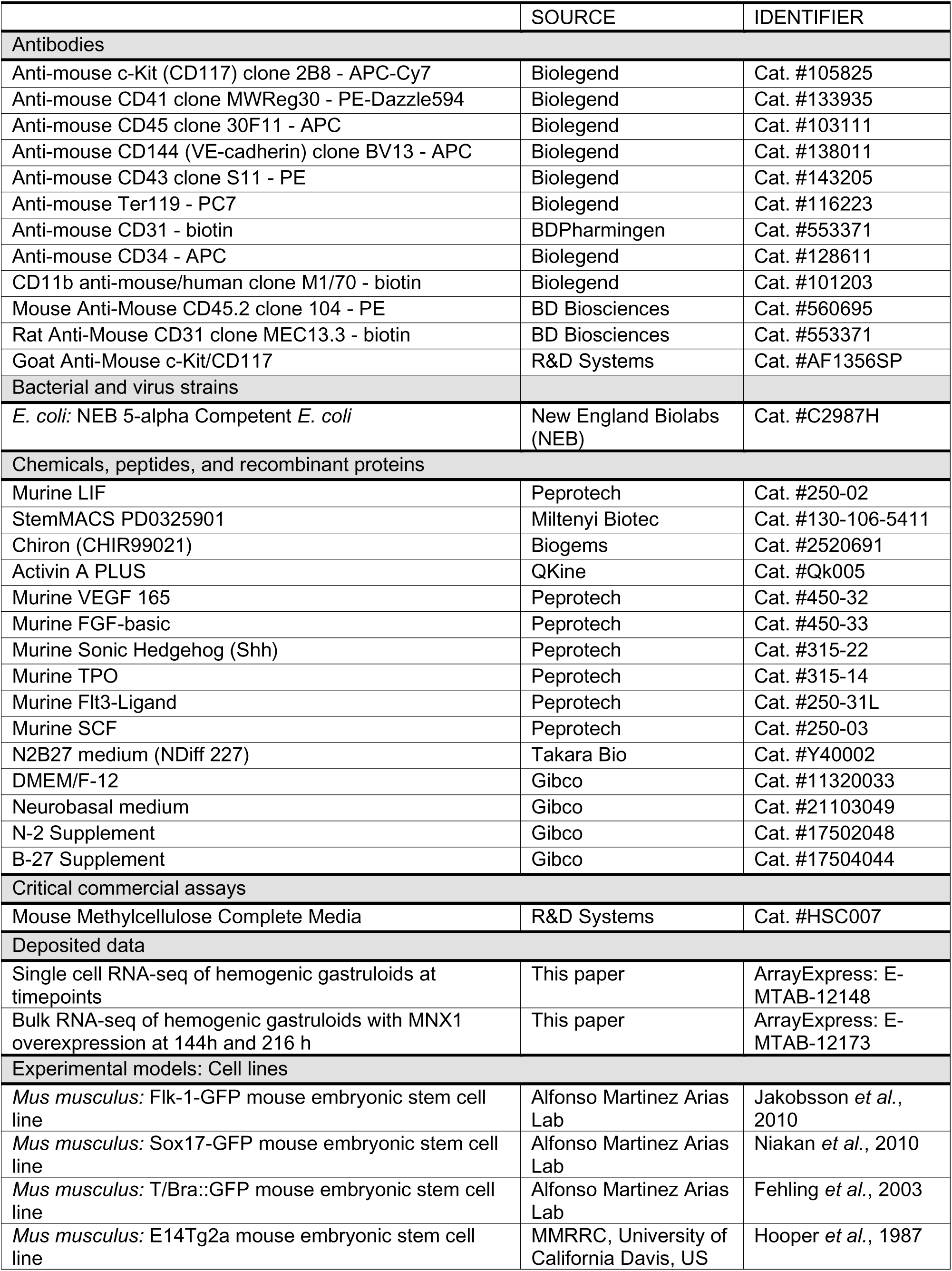
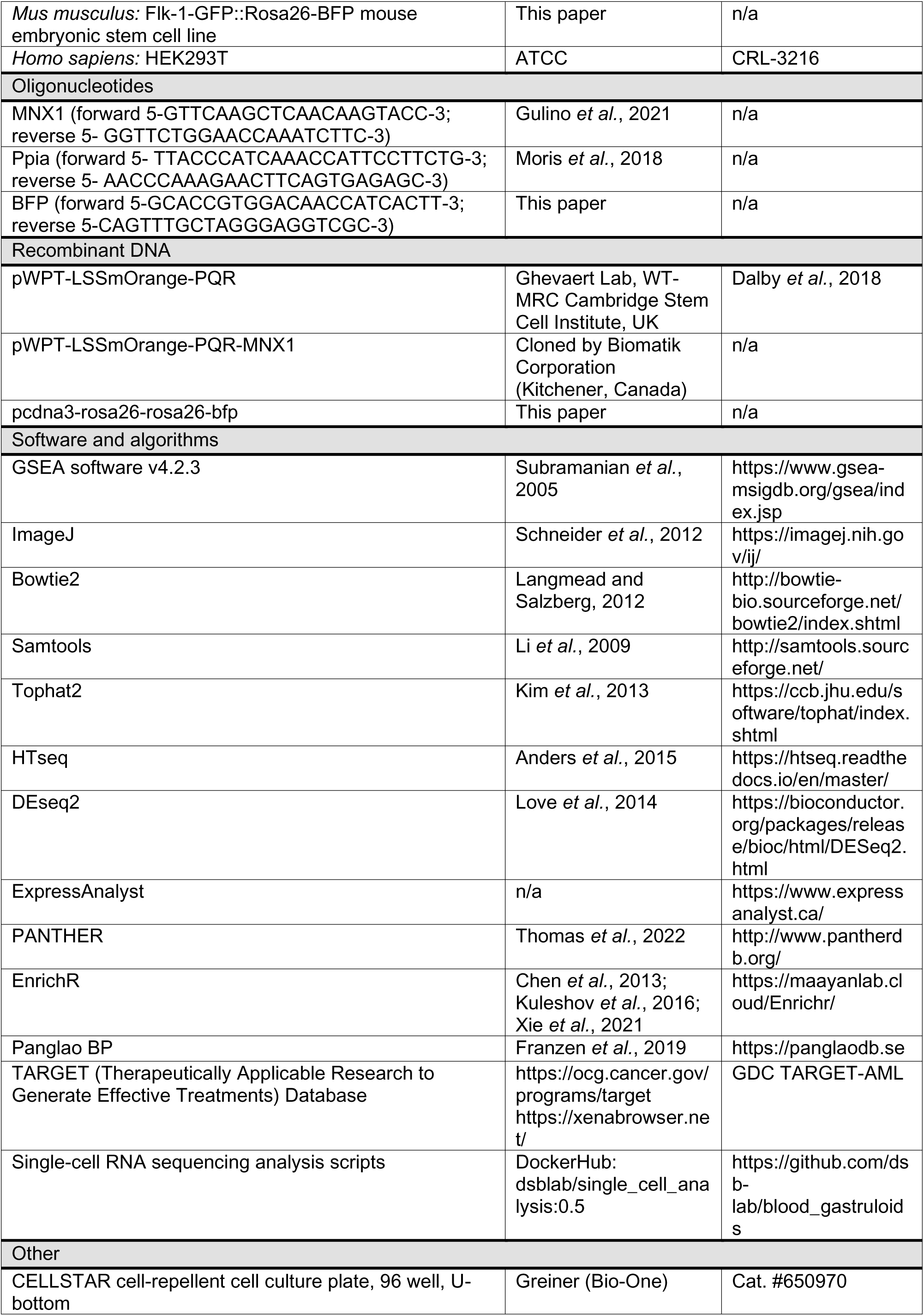

### METHODS

#### Cell culture

Flk-1-GFP (Jakobsson *et al.,* 2010), Flk-1-GFP::Rosa26-BFP, Sox17-GFP (Niakan *et al.,* 2010), T/Bra::GFP (Fehling *et al.,* 2003), and E14Tg2A (Hooper *et al.,* 1987) mouse embryonic stem cells (mES) lines were cultured in ES+LIF medium in gelatinized flasks (0.1% gelatin) with daily medium change, as previously described (Turner *et al.,* 2017). The ES+LIF medium contained 500 ml Glasgow MEM BHK-21 (Gibco), 50 ml of fetal bovine serum (FBS, Embryonic Stem Cells tested, biosera, Nuaillé, France), 5 ml GlutaMAX supplement (Gibco), 5 ml MEM Non-Essential Amino Acids (Gibco), 5 ml Sodium pyruvate solution (100 mM, Gibco), and 1 ml 2-Mercaptoethanol (50mM, Gibco). Murine LIF (Peprotech) was added at 1000 U/ml. HEK293T cells were grown in Dulbecco’s modified Eagle medium (Gibco) supplemented with 10% FBS (Gibco) and 1% penicillin/streptomycin (Gibco). All cultures were kept at 37°C and 5% CO_2_.

#### Hemogenic gastruloids assembly, culture, and dissociation

mES were maintained in ES+LIF medium and transferred to 2i+LIF (containing Chiron and MEK inhibitor PD03) for 24 hours prior to the assembly into gastruloids. 250 untransduced mES cells (or 400 for GX-MNX1 and GX-EV) were seeded in each well of a U-bottom, cell-repellent 96-well plate (Greiner Bio-One, Stonehouse, UK) in 40 μl of N2B27 medium [Takara Bio or home-made as Cold Spring Harbour Protocols ‘N2B27 Medium’ (2017)]. The plate was centrifuged at 750 rpm for 2 minutes to promote deposition and aggregation of the cells and was then incubated at 37°C, 5% CO2 for 48 hours. After 48 hours, 150 μl of N2B27 medium supplemented with 100 ng/ml Activin A (QKine, Cambridge, UK) and 3 μM chiron (Peprotech) was added to each well. At 72 hours, 150 μl of medium were removed, without disrupting the gastruloids in the wells. 100 μl of N2B27 with 5 ng/ml of Vegf and Fgf2 each (Peprotech) were added to each well. From 72 h to 144 h, each day 100 μl of medium were removed and replaced with N2B27 + Vegf + Fgf2. At 144 h, the medium was further supplemented with Shh at 20 ng/ml. From 168 h to 216 h, the medium was N2B27 + 5 ng/ml Vegf, plus 20 ng/ml mTpo, 100 ng/ml mFlt3l, and 100 ng/ml mScf (Peprotech). To dissociate cells from the gastruloid structures, medium was removed and individual gastruloids were collected using a pipette and precipitated at the bottom of a microcentrifuge tube. The remaining medium was aspirated and the bulk of gastruloids was washed in PBS. 50 μl of TrypLE express was added to pelleted gastruloids to be incubated at 37°C for 2 minutes to dissociate cells.

#### Animals and implantation

Female athymic nude-Foxn1^nu^ (*nu/nu*) mice (Envigo, Bresso, Italy) were housed under pathogen-free conditions. In accordance with the “3Rs policy”, experiments were reviewed and approved by the Animal Welfare Body (OPBA) of IRCCS Ospedale Policlinico San Martino and by the Italian Ministry of Health (n. 883/2020-PR).

Intact haemGx (3 per mouse in 10 μL of PBS) were inoculated in the adrenal gland of five-week-old mice. Briefly, mice were anesthetized with a mixture of xylazine (ROMPUN, Bayer) and ketamine (LOBOTOR, Acme S.r.l.) and injected with haemGx, after laparotomy, in the capsule of the left adrenal gland, as previously described (Brignole *et al.,* 2023; Pastorino *et al.,* 2003). All mice survived surgery. Control mice (CTR) received vehicle only. Mice body weight and general physical conditions were recorded daily. One month after haemGx inoculation, mice were sacrificed and spleen and bone marrow (BM) from femurs collected. BM cells and spleens, mechanically reduced to cell suspension, were stored at −80°C until use. Adrenal glands were paraffin-embedded for subsequent IHC studies.

#### Methylcellulose colony forming assays (CFC)

Disassembled gastruloids cells were plated on Mouse Methylcellulose Complete Media (R&D Systems). Cells were first suspended in Iscove’s Modified Dulbecco’s Medium (IMDM, Gibco) with 20% FBS (Gibco) before addition to the methylcellulose medium. Cells were plated in duplicate 35 mm dishes with 2×10^5^ cells/plate. Plates were incubated at 37°C and 5% CO2 for 10 days, when colonies were scored. For serial replating experiments, cells in methylcellulose were collected and washed in phosphate buffer saline (PBS) to achieve single-cell suspensions and replated as described above. Dissociated gastruloid cells from CFC were collected as cell suspensions in PBS and centrifuged onto slides (Shandon Cytospin 2 Cytocentrifuge) at 350rpm for 5-7min, with a PBS-only pre-spin at 1000rpm, 2min. Slides were stained in Giemsa-Wright stain for 30 seconds followed by addition of phosphate buffer pH 6.5 for 5 min.

#### Immunofluorescence staining

Immunostaining of whole gastruloids was performed as described before (Baillie-Johnson *et al*., 2015). Briefly, gastruloids were fixed in 4% paraformaldehyde (PFA) dissolved in PBS for 4 hours at 4°C on orbital shaking and permeabilised in PBSFT (10% FBS and 0.2% Triton X-100), followed by one hour blocking in PBSFT at 4°C on orbital shaking. Antibody dilutions were made in PBSFT at 1:200 for primary and 1:500 for secondary antibodies. Antibody incubations were performed overnight at 4°C on orbital shaking, and subjected to optical clearing overnight in ScaleS4 clearing solution. Individual gastruloids were then mounted on glass coverslips by pipetting as droplets in ScaleS4 and DAPI nuclear stain.

#### Histology and H&E staining

Samples were fixed with 10% formalin at room temperature for 48 hours followed by an ethanol series of ethanol and paraffin; ethanol 70% (90 minutes), ethanol 80% (120 minutes), ethanol 95% (180 minutes), 3 times ethanol 100% of 120, 120 and 180 minutes, 50% ethanol/paraffin (3×120 minutes) and 3 times paraffin (60, 90 and 120 minutes) before embedding in histological cassettes. Blocks were sectioned with a HistoCore BIOCUT Microtome (Leica) at 10 mm and slides were stained with Haematoxylin/Eosin.

#### Imaging

Images of cultured gastruloids and CFC plates were captured using the Cytation 5 Cell Imaging Multi-Mode Reader (Biotek) plate reader using bright field and FITC channels. ImageJ (Schneider *et al*., 2012) was used for gastruloid size quantification. Confocal microscopy was performed on LSM700 on a Zeiss Axiovert 200 M with Zeiss EC Plan-Neofluar 10x/0.30 M27 and Zeiss LD Plan-Neofluar 20x/0.4 M27 objective lens. Tissue sections were imaged with a THUNDER Imager 3D microscope system (Leica).

#### Lentiviral vector packaging and transduction

The lentiviral overexpression vector pWPT-LSSmOrange-PQR was used to clone the *MNX1* gene cDNA. The viral packaging vectors pCMV and pMD2.G, described in Pina *et al*. (2008), were used to assemble lentiviral particles using HEK293T cells via transfection using TurboFect Reagent (Invitrogen). Transduction of mES cells was performed overnight by addition of lentivirus to culture medium and washed the following day (Moris *et al.,* 2018). Transduced cells were sorted for positivity to LSSmOrange.

#### Flow cytometry

Surface cell marker analysis was performed by staining using the antibodies listed in Key Resources Table. Disassembled gastruloids cells were resuspended in PBS, 2% FBS and 0.5 mM EDTA and stained at a dilution of 1:100 for primary antibodies for 20 minutes at 4°C. When indicated, streptavidin was added at a dilution of 1:200. Analysis was performed on ACEA Novocyte (Agilent) or AttuneNxT (Thermo) analyzers, using the respective software packages. Cell sorting was performed using a CS&T calibrated BDFACS Aria III system (488nm 40 mW, 633nm 20 mW, 405nm 30 mW, and 561nm 50 mW), set with the 100μm nozzle at 20PSI and a 4-way purity mask or, in the case of haemGx adrenal implants, a Beckman Coulter CytoFlex SRT (488nm 50 mW, 633nm 100 mW, 405nm 90 mW, and 561nm 30 mW), set with 100μm nozzle at 15PSI, also using 4-way purity. Single-cell deposition in 96-well plates was performed using single-cell sorting mode. Intact cells were gated on FSC-A vs SSC-A plot, followed by doublet exclusion on FSC-A vs FSC-H and SSC-A vs SSC-H, prior to gating on fluorescent parameters for the markers described in the results.

#### Single cell RNA sequencing of time-resolved hemogenic gastruloids

Gastruloids were collected at different timepoints of the protocol, disassembled and FACS deposited into 96-well plates, either as unsorted (global) or sorted by CD45, CD41 and C-Kit/ScaI markers (sorted). RNA from the cells were extracted from single cells using Smart-seq2 technology at 500Kb and depth of 151 bases. Sequencing reads were quality-checked using FastQC (v0.11.9). We trimmed the samples using trimGalore! (0.6.6) with a cutoff of 30, clipping 15 base pairs and retaining reads of more than 100 bases. Alignment was performed on STAR (2.7.8a) and Mus musculus annotations from GENCODE vM26. Aligned BAM files were annotated using featureCounts (v2.0.1) and count matrices were computed in python by directly accessing the BAM files with gtfparse packages and collapsing lanes.

For quality control, based on the histogram of counts and multimodality distributions, we set a minimum count threshold of 200000 counts, a minimum threshold of 1000 expressed genes, and a maximum threshold of 20% mitochondrial fraction per cell. We performed the same procedure of the second dataset with a more restrictive threshold of 400000 counts and similar expressed genes and mitochondrial thresholds. Cells that did not pass the quality control metrics were omitted from analysis. We normalized the cells to the mean count number per dataset and applied a plus-one-log transformation of the data before proceeding to the downstream analysis.

Dimensionality reduction was performed on feature selection of the gene space using the function scanpy.highly_varying_genes with default parameters. Selection of principal components in principal component analysis (PCA) was performed by heuristic elbow method. Nearest neighbor analysis was constructed by KNN graphs using a correlation metric and 10 nearest neighbors. Data was projected for low dimensional visualization using the UMAP algorithm with default parameters as implemented in scanpy.tl.umap. We used the leiden algorithm as implemented in scanpy.tl.leiden to partition the data into clusters. In order to assess the election of the resolution parameter we used Newman-Girvan modularity as a metric of clustering quality. Differential expressed genes were computed comparing each cluster against the rest using the Wilcoxon test with Benjamini-Hochberg correction.

#### Annotation and projection to additional datasets

Raw counts matrices were downloaded and processed following the same pipeline as described above for scRNA-seq of gastruloids to generate UMAP and clustering. We implemented the scmap algorithm to compare our scRNA sequencing of gastruloids with the available datasets. We reduced the dimensionality of the space by selecting highly varying genes from the annotated dataset. Then, we constructed a KNN classifier with correlation metric and computed the nearest neighbors of the target data. If neighbors with correlation metrics below 0.7 default standards, the projected cells were not projected onto any cell from the annotated dataset. To visualize the cells over the UMAP plots of the other datasets, we constructed a KNN regressor with three neighbors and a correlation metric. Confusion matrices were used to visualize the overlap between gastruloid clusters and the embryo dataset assigned clusters using scmap.

#### Bulk RNA sequencing

Total RNA was extracted from disassembled gastruloid cells at 216 hours. Sequencing was performed on NovaSeq PE150 platform, at 20M paired-end reads per sample. Tophat2 with Bowtie2 were used to map paired-end reads to the reference Mus musculus genome build GRCm39. GENCODE Release M30 (Frankish *et al.,* 2019) was used as the reference mouse genome annotation. Aligned reads were filtered by quality using samtools (Li, H. *et al.,* 2009) with a minimum threshold set at 30 (q30). Transcript assembly and quantification was achieved using htseq (Putri *et al*., 2022). Differential expression between sample and control was performed by collapsing technical replicates for each condition using Deseq2 (Love, Huber and Anders, 2014) in R environment (Deseq2 library v 1.32.0). The differential expression was expressed in the form of log2 fold change and filtered by false discovery rate (FDR) of 0.05.

#### Gene list enrichment analyses

Gene ontology (GO) analysis was performed in ExpressAnalyst (available at www.expressanalyst.ca) using the PANTHER or GO Biological Process (BP) repository. GO terms and pathways were filtered by false discovery rate (FDR) with a cut-off of ≤ 0.1 for meaningful association. EnrichR (Xie *et al.,* 2021; Kuleshov *et al.,* 2016; Chen, E. Y. *et al.,* 2013) was used for cell type analysis using the Panglao DB (Franzén, Gan and Björkegren, 2019) databases using a FDR threshold of ≤ 0.1 or p value ≤ 0.01, where specified, as well as transcription factor (TF) binding site enrichment using the ENCODE and ChEA Consensus TFs from ChIP database.

#### Gene Set Enrichment Analysis (GSEA)

Custom gene signatures were used as gene sets for GSEA analysis (Subramanian *et al.,* 2005) on the GSEA software v4.2.3 on RNA sequencing expression values in counts units. GSEA was ran in 10000 permutations on gene set using the weighted Signal2Noise metric. Enrichment metrics are shown as normalized enrichment score (NES) and filtered by FDR ≤ 0.05. Leading edge genes (LEGs) are genes with a “Yes” values for core enrichment. For AML patient analysis, clinical phenotype and expression data (in counts units) were extracted from the GDC TARGET-AML cohorts in the Therapeutically Applicable Research to Generate Effective Treatments project (TARGET, https://ocg.cancer.gov/programs/target), downloaded from the University of California Santa Cruz (UCSC) Xena public repository (last accessed 31st August 2022). Patient samples were selected according to the reported karyotype to include t(7;12), inv(16), MLL, normal karyotype, and t(8;21). GSEA was performed comparing RNA sequencing counts of t(7;12) samples against pooled AML subtypes (inv(16), MLL, normal karyotype and t(8;21)) as “other AML”.

#### Real-time polymerase chain reaction (qPCR)

Extracted RNA was reverse-transcribed into complementary DNA (cDNA) using High-Capacity RNA-to-cDNA Kit (Applied Biosystems). QPCR was performed using FastGene 2x IC Green Universal qPCR Mix (Nippon Genetics, Duren, Germany) using primers for human *MNX1* (forward 5-GTTCAAGCTCAACAAGTACC-3; reverse 5-GGTTCTGGAACCAAATCTTC-3) (Gulino *et al*., 2021) and *Ppia* for endogenous control (forward 5-TTACCCATCAAACCATTCCTTCTG-3; reverse 5-AACCCAAAGAACTTCAGTGAGAGC-3) (Moris *et al.,* 2018), or Taqman gene expression assays (Mm00501741_m1 *Myb*; Mm00439364_m1 *Hoxa9*; Mm00723157_m1 *Hlf*; Mm01213404_m1 *Runx1*; Mm00444619_m1 *Dll4*; Mm00491303_m1 *Mecom*; Mm02342430_g1 *Ppia*) using Taqman Gene Expression Mastermix (Applied Biosystems). Differential gene expression was calculated using the delta delta Ct (ΔΔCt) method.

#### Polymerase chain reaction (PCR)

Genomic DNA was extracted using Monarch Genomic DNA Purification Kit (New England Biolabs, Hitchin, UK) using manufacturer’s instructions. PCR on genomic DNA was performed using Phire PCR Master Mix (Thermo Scientific) to amplify the BFP locus using forward primer 5’-GCACCGTGGACAACCATCACTT-3’ and reverse primer 5’-CAGTTTGCTAGGGAGGTCGC-3’.

#### Statistical analysis

Experiments were performed at least in triplicates, unless specified otherwise. Data are plotted to include standard deviation (+/−SD) between replicates. Statistical significance was set at a threshold of p value < 0.05. Statistical analysis was performed in R environment (version 4.1.3) or using GraphPad Prism 8.0 software and is detailed in respective figure legends.

## Data availability

Raw data as well as processed count matrices and post-processed files from single-cell RNA-seq for the time-resolved data is available at E-MTAB-12148. Bulk RNA-seq of MNX1 overexpressing gastruloids is available at Array Express with accession code E-MTAB-12173. The post-processing was performed in Python on DockerHub: dsblab/single_cell_analysis:0.5. Scripts are available in https://github.com/dsb-lab/blood_gastruloids and Zenodo (https://doi.org/10.5281/zenodo.7053423). The results published here are partly based upon data generated by the Therapeutically Applicable Research to Generate Effective Treatments (TARGET) (https://ocg.cancer.gov/programs/target) initiative, of the Acute Myeloid Leukemia (AML) cohort GDC TARGET-AML. The data used for this analysis are available at https://portal.gdc.cancer.gov/projects and https://xenabrowser.net/.

## ACKNOWLEDGEMENTS

This project was funded by a start-up grant and a BRIEF award from Brunel University London, a Little Princess Trust (LPT) Project Grant (CCLGA 2023 22 Pina) and a National Centre for the Replacement, Refinement and Reduction of Animals in Research (NC3Rs) project grant (NC/Z500677/1) to CP, by an Agencia Estatal de Investigación grant (PLEC2021-007518) to AB and AMA, and by an ERC Advanced Grant (MiniEmbryoBlueprint 834580) to AMA. LPT research is funded in partnership with CCLG through the CCLG Charity Research Network. DR received funding from the Royal Society of Biology (MRSB Travel Grant). GTC was funded by grant FPU18/05091 from the Spanish Ministry of Universities. AJ is funded by a Lady Tata Memorial Trust International Scholarship (2022). SvdB was funded by an EMBO Postdoctoral Fellowship (ALTF 195-2021) and is in receipt of a HFSP Postdoctoral Fellowship (LT0047/2022-L). Work in the CP lab was also funded by a KKLF Intermediate Fellowship (KL888), a Leuka John Goldman Fellowship for Future Science (2017-2019), and a Wellcome Trust / ISSF Bridge Funding award at the University of Cambridge (2019). JGO acknowledges financial support from the Spanish Ministry of Science and Innovation and FEDER (grant PGC2018-101251-B-I00), by the Maria de Maeztu Programme for Units of Excellence in R&D (grant CEX2018-000792-M), and by the Generalitat de Catalunya (ICREA Academia programme). MP acknowledges funding from the Italian Ministry of Health (Ricerca Finalizzata 5 per mille and Ricerca Corrente to MP). Library preparation and next-generation sequencing for single-cell RNA-seq analysis were performed by the Single Cell Genomics Group at the National Centre for Genomic Analysis – Centre for Genomic Regulation (CNAG-CRG), Barcelona. The Authors wish to acknowledge Tina Balayo, Ana Filipa Domingues, Shikha Gupta, Oliver Davies, Kristen Place and Remisha Gurung’s technical support at different stages of the project.

## AUTHOR CONTRIBUTIONS

Conceptualization: CP, AMA; Methodology: CWS, DR, GTC, FP, SvdB, MC,MP, KRK, VH-H, AB, JGO, AMA, CP; Software: GTC, JGO; Validation: CWS, DR, AJ, YC, SvdB, CP; Investigation: CWS, DR, GTC, FP, AJ, YC, LD, CB, G-AI, CP; Formal Analysis: GTC, DR, JGO, CP; Resources: GTC, JC, MP, JGO, AMA; Data curation: DR, GTC, JGO; Writing – Original Draft: CP, DR, GTC; Writing – Review and editing: CP, DR, GTC, AB, JGO, AMA; Visualisation: DR, GTC, CWS, CP; Supervision: CP; Project administration: CP, AMA, JGO; Funding acquisition: CP, AMA, AB.

## SUPPLEMENTAL INFORMATION

**Figure 1 Supplement 1 Optimization and reproducibility of the haemGx protocol. (A)** Imaging of a 120h-haemGx with and without the addition of Activin A in conjunction with Chiron 99021 (CHIR) to achieve polarized expression of *Flk-1-GFP*; scale bar: 100mm. **(B)** Titration of Activin A dosage (from 25 ng/ml to 150 ng/ml) required for *Flk-1-GFP* expression activation (green). **(C)** Representative flow cytometry plots of %CD45^+^ cells at 216h without and without addition of SCF, Flt3L and TPO (SFT) from 168h; representative plots. **(D)** Quantification of %CD45^+^ cells by flow cytometry analysis of haemGx cultured with or without SFT, between 192-216h and 168-216h. Mean ± SD of n=7-8 independent experiments; Tukey’s multiple testing with significant q-value<0.05. **(E)** Quantification of %CD45^+^ cells at 216h by flow cytometry analysis of haemGx cultured in VEGF+FGF or VEGF-only during the 168-216h period. Mean ± SD of n=7 independent experiments; unpaired t-test; ns, not significant. **(F)** Flow cytometry monitoring of C-Kit, CD41, and CD45 markers in haemGx established from E14 and *Flk-1-GFP* mES lines; n=3-4 replicate experiments (n=1, CD45^+^ 120h). Mixed effects model with Sidak’s multiple comparison test; significant adjusted p-value <0.05. **(G)** Representative flow cytometry plots of CD41^+^ and C-Kit^+^ cells in E14 and *Flk-1-GFP* mES cell-initiated haemGx. **(H)** Representative flow cytometry plots of CD45^+^ cells in E14 and *Flk-1-GFP* mES cell-initiated haemGx. **(I)** Quantification of %CD45^+^ cells in *Flk-1-GFP*, *Sox17-GFP*, and *TBra-GFP* mES cell-initiated haemGx; n=5, mean ± SD. Welch’s t-test Sox17-GFP vs. Flk1-GFP, p=0.5254; Tbra-GFP vs. Flk1-GFP, p=0.3782.

**Figure 2 Supplement 1 Multi-color flow cytometry detection of surface markers in haemGx. (A)** Representative flow cytometry plots of CD31 and CD34 staining of haemGx at 144h and 216h gated on Flk-1-GFP^+^. **(B)** Quantification of Flk-1-GFP^+^ haemGx populations at 144h and 216h expressing CD34 and CD31. Crossbar shows mean of 4 replicate experiments with individual data points shown; Welch’s t-test; p value significant < 0.05. **(C-E)** Unstained controls (C-D) and fluorescence minus one (FMO) controls (E) relative to flow cytometry detection of multi-color of surface markers in Figure 2. **(F)** Representative flow cytometry plot of haemGx of CD45^+^CD41lo fractions co-expressing C-Kit and Cd144 (Ve-Cadherin).

**Figure 2 Supplement 2 Characterization of hematopoietic output from haemGx. (A)** Representative image of cytospins of dissociated haemGx at 216h stained with Giemsa-Wright’s stain. Annotated are cells in the monocytic (dashed open arrow), granulocytic (solid open arrow), megakaryocytic (solid arrow) and erythroid (asterisk) lineages; arrowheads indicate cells with a non-specific blast-like morphology. 40x magnification, scale bar = 100 µm. **(B)** Representative flow cytometry plots of CD41 and CD45 expressing populations in haemGx treated with 0.5 μM EZH2 inhibitor GSK126 or with 0.05% DMSO (control) at 144h and 216h. **(C)** Quantification of flow cytometry analysis in (B) showing the proportion of CD41^+^ and CD45^+^CD41^lo^ populations in GSK126-treated haemGx. Welch’s t-test; p value significant < 0.05.

**Figure 3 Supplement 1 Single-cell-RNAseq (scRNAseq) analysis of haemGx identifies time-dependent signatures of endothelial, hemogenic and stromal cells. (A)** Summary of plating strategy for scRNA-seq analysis of gastruloids at 120h, 144h, 168h, 192h, and 216h without selection of surface markers (‘all’) or sorted as C-Kit/Sca1+ (green shading), CD41+ (blue), or CD45+ (orange) cells; P = plate. **(B)** ScRNA-seq quality control measures (quantification of mapped reads, individual genes and mapping of reads on to mitochondrial (mt)DNA. **(C)** Mapped read and gene counts for individual Smart-seq2 libraries (plate, P), biological replicate (repl) populations of single cells, and read and map distributions across unsorted and cell surface phenotype-sorted cells. **(D)** UMAP projection of all sequenced cells colored by annotated clusters (left), by sorting markers C-Kit/ScaI, CD41, and CD45, or unfractionated cells (‘whole’ corresponding to ‘all’ in panel A) (center), and by haemGx culture time (right). **(E)** Cell type enrichment analysis of cluster classifier genes, using the PanglaoDB repository (Franzén, Gan and Björkegren, 2019); classifier genes are obtained by differential expression in comparison to all other clusters. Represented are clusters 1, 3 and 9, which are characteristic of timepoints 168-216h, and capture stromal cells putatively relevant for hemogenic support, including autonomic neurons (cluster 9). The statistical power of representation of individual cell types is expressed as the EnrichR combined score with a p-value threshold of < 0.05.

**Figure 3 Supplement 2 Single-cell-RNAseq (scRNAseq) analysis of haemGx. (A)** Heatmap representation of the expression of endothelial and hematopoietic marker genes in individual haemGx CD41^+^ and CD45^+^ cells sorted at 192 and 216h. Cells and genes ordered by unsupervised hierarchical clustering. **(B)** Violin plots quantifying the expression of arterial markers in C-Kit+ fraction from clusters in Fig. 3A comparing 144h and 192h timepoints. Wilcoxon test; significant p value<0.05. **(C)** Summary representation of the proportion of hGx cells expressing endothelial (endo), erythroid/myeloid (Ery+My), myeloid/lymphoid (My+Ly) and HSPC gene signatures. Endo: *Kdr*; Ery+My: *Epor*±*Gata1*±*Klf1*±*Hbb*±*Hba*±*Hbg* + *Spi1*±*Mpo*±*Anpep*±*Csf1/2/3r*); My+Ly: *Spi1*±*Csf1/2a/3r* + *Ikzf1*±*Ighm/d*±*Igk*); HSPC: *Ptprc* + *Myb*).

**Figure 4 Supplement 1 Analysis of mouse embryonic scRNA-seq datasets for transcriptional similarities of haemGx outputs. (A-C)** UMAPs of mouse single-cell RNA-seq datasets annotated as in respective publications: (A) arterial and haemogenic specification in the para-splanchnopleura (pSP) and AGM region between E8.0 and E11 (Hou et al. 2020); (B) YS, AGM and FL progenitors and the AGM EHT (Zhu et al., 2020); (C) HSC emergence from the dorsal aorta HE (Thambyrajah et al., 2024).

**Figure 4 Supplement 2 Detection of engraftment potential of late-stage haemGx in adrenal glands of Nude mice. (A)** Map of the *Rosa26-BFP* construct used to target *Flk1-GFP* mES cells employed in implant assays. Primers used for gDNA PCR analysis are indicated. **(B)** PCR detection limit of the BFP amplicon using the primers in (A), in a serial dilution of *Rosa26-BFP*::*Flk1-GFP* mES gDNA into human leukemia cell line K562 gDNA; total 100ng DNA per lane. **(C)** Uncropped image of agarose gel electrophoresis of BFP detection shown in Fig. 4C. **(D)** Representative flow cytometry plot of the spleen of immunodeficient mouse recipients, showing myelo-lymphoid (CD45^+^Ter119^-^) and erythroid (Ter119^+^CD45^+/−^) cell fractions at 4 weeks post implantation. **(E)** PCR analysis of *Rosa26-BFP*::*Flk1-GFP* gDNA in the cell fractions in (D) of implanted animal #15 at 4 weeks post-implantation. PCR run in triplicate material from each sample; positive and NTC controls as per Fig. 4C. **(F-G)** Representative flow cytometry interrogation of BFP detection within lineage compartments in the bone marrow (BM) of control and implanted recipient animal #2 (F) and animal #1 (G) at 8 weeks after hGx implantation. **(H)** Histology of implanted and control adrenal glands; paraffin sections stained with hematoxylin & eosin (H&E). Inserts in implanted animal highlight neural (orange) and hematopoietic and renal tubular epithelium (green); inserts in control animal highlight normal cortical (blue) and medullar (red) adrenal architecture, absent in implanted adrenal glands.

**Figure 5 Supplement 1 MNX1 overexpression promotes hemogenic specification in haemGx. (A)** Cell type enrichment analysis in MNX1-r AML transcriptomes, compared to normal paediatric bone marrow (BM) or other paediatric AML from the TARGET database. GSEA used representative cell type gene sets from the 2021 DB database; bubble plot shows NES scores (color gradient) and statistical significance (–log10(FDR), bubble size). **(B)** Quantitative (q)RT-PCR analysis of *MNX1* overexpression in 216h-haemGx. Gene expression fold change calculated by normalization to HPRT1. Mean ± SD of 3 replicates; 2-tailed t-test, p< 0.001 (**). **(C)** Cell counts of disassembled haemGx at 216h. Mean ± SD of 3 replicate experiments; 2-tailed t-test, p<0.001 (**). **(D)** Proportion of EV and MNX1 haemGx exhibiting spontaneous unifocal contractility at 192h, observed in commercial N2B27 medium (see Experimental Procedures); mean ± SD of 3 replicate experiments; 2-tailed t-test, p<0.001 (**). **(E-F)** Flow cytometry analysis of specification of **(E)** hemato-endothelial – C-Kit^+^ and VE-Cadherin^+^ – and **(F)** hematopoietic progenitor – CD41^+^ and CD45^+^ – cells over a 120-216h timecourse of EV and MNX1 haemGx cultures; representative plots.

**Figure 5 Supplement 2 Transcriptional analysis of haemGx with MNX1 overexpression. (A)** Expression of human *MNX1* and murine *Mnx1* genes in FPKM units from RNA-seq of MNX1 and EV haemGx. **(B)** UMAP of time-resolved global clustering of scRNA-seq of haemGx showing the enriched clusters in GX-MNX1 at 144 hr (orange boxes) and 216 hr (blue boxes), and MNX1-*r* AML (black boxes) or MLL-AML (green boxes) determined by GSEA. **(C)** Bubble plot of GSEA NES values and statistical significance by – log10(FDR) of enrichments in specific time clusters and corresponding global clusters in GX-MNX1 at 144 hr and 216 hr compared to respective GX-EV, and MNX1-*r* or MLL AML samples compared to other pediatric AML.

**Figure 6 Supplement 1 Characterization of MNX1-overexpressing replating cells from haemGx. (A)** Representative flow cytometry plot of CD31 and C-Kit staining of CFC plating of GX-MNX1 (left) and quantification of sustained expression of CD31^+^CKit^+^ cells through replatings. **(B)** Representative flow cytometry plots of serially re-plating MNX1 cells from 144h and 216h-gastruloids (plate 5 cells) stained with Ter119 and CD45.

**Figure 7 Supplement 1 Transcriptional analyses of haemGx with *MNX1* overexpression compared to patient signatures and current *in vivo* model. (A)** Venn diagram showing the intersection of leading-edge genes enriched in *MNX1*-r patient vs MLL, core-binding factors (CBF), or other pediatric AML from GSEA analysis using the K14 signature. **(B)** Cell type enrichment analysis in *MNX1*-r AML transcriptomes, compared to normal paediatric bone marrow (BM) or other paediatric AML from the TARGET database, with comparison with transcriptomes of haemGx and CFCs at from MNX1 vs EV conditions at 144h and 216h. GSEA used representative cell type gene sets from the Panglao DB 2021database; bubble plot shows NES scores (color gradient) and statistical significance (–log10(FDR), bubble size). **(C)** Representative Giemsa-Wright stained dissociated CFC replating MNX1 haemGx cells from 144h; purple arrows, undifferentiated blasts; solid black arrow, mast cell precursors; dashed black arrow, mast cells. **(D)** Transcriptional re-analysis of Waraky et al (2024) in vivo model of MNX1-OE leukemia from transplanted fetal liver (FL) cells. Heatmap comparing the expression levels of all DEG between all conditions, i.e. FL control (FL Ctrl) vs FL with MNX1-OE (FL MNX1) in liquid culture, and bone marrow (BM) from mice transplanted with FL Ctrl (BM TX Ctrl) vs leukemic animals transplanted with FL MNX1 cells (BM TX MNX1). Hierarchical clustering by Ward D method on Euclidean distances identifies 12 clusters (k). Clusters k1, k2, k5, k6 and k9 are specifically up-regulated in BM TX MNX1 and putatively represent MNX1-OE leukemia programs. Analysis matches Fig. 7B for MNX1-OE haemGx. **(E)** GSEA plots of significant results (by p value < 0.05) for gene set extracted from k1, k2, k5, k6 and k9, individually or combined, from (D) against MNX1-r patients RNA-seq counts vs MLL, core-binding factors (CBF), or other pediatric AML. FDR values are indicated for each enrichment point. Analysis matches Fig. 7C for MNX1-OE haemGx. **(F)** Cell type enrichment analysis in MNX1-r AML transcriptomes, compared to normal pediatric bone marrow (BM) or other pediatric AML from the TARGET database. GSEA used representative cell type gene sets from the Panglao DB 2021 database; bubble plot shows NES scores (color gradient) and statistical significance (–log10(FDR), bubble size). Analysis matches Fig. S6E for MNX1-OE haemGx.

Supplemental Files S1-S3 (Excel files):

Supplemental File S1: Cluster classifier gene lists for each cluster by differential gene expression analysis of scRNA-seq of haemGx (all conditions), obtained by Wilcoxon rank test of each cluster against all other clusters.

Supplemental File S2: Cell type enrichment analysis of cluster classifier genes, showing the inferred identity of clusters of scRNA-seq of haemGx (all conditions). Classifier genes (in Supplemental File S1) were obtained by differential expression in comparison to all other clusters and subjected to enrichment analysis using the PanglaoDB repository (Franzén, Gan and Björkegren, 2019).

Supplemental File S3: Summary of projections of scRNA-seq of haemGx cells onto available datasets by Hou et al. (2020), Zhu et al. (2020), and Thambyrajah et al. (2024), showing the identity of each haemGx cell projecting to annotated clusters of the respective studies.

Supplemental File S4: List of differentially expressed genes between haemGx-MNX1 and haemGx-MNX1 at 144h and 216h compared to empty vector (EV) control.

