## Supplementary Figures for "Dissecting infant leukemia developmental origins with a hemogenic gastruloid model"

[1]College of Health, Medicine and Life Sciences, [2]Centre for Genome Engineering and Maintenance, Brunel University London, UK; [3]Department of Genetics, University of Cambridge, UK; [4]Department of Medicine and Life Sciences, Universitat Pompeu Fabra, Spain; [5]Laboratory of Experimental Therapies in Oncology, IRCCS Istituto Giannina Gaslini, Genova, Italy; [6]Program in Cancer Research, Hospital de Mar Research Institute, CIBERONC, Barcelona, Spain; [7]Josep Carreras Leukemia Research Institute, Barcelona, Spain; [8]Animal Facility, IRCCS Policlinico San Martino, Genova, Italy; [9]Department of Pathology, University of Cambridge, UK; [10]Centre for Haemato-Oncology, Barts Cancer Institute, Queen Mary University of London, UK; [11]Centre for In Vivo Modelling, Institute of Cancer Research, Sutton, UK.

[12]Current address: Sanquin Research, Landsteiner Laboratory, Amsterdam University Medical Center, Amsterdam, The Netherlands

[13]Current address: Faculty of Biology, Medicine and Health, University of Manchester, Manchester, UK

[14]Current address: Division of Infection and Immunity, Cardiff University School of Medicine, Cardiff, Henry Wellcome building, CF14 4XN, UK

[15]Current address: Sanquin Research, Amsterdam, The Netherlands

*These authors contributed equally to this work

#### Supplementary Files

**Supplemental File S1**: Cluster classifier gene lists for each cluster by differential gene expression analysis of scRNA-seq of haemGx (all conditions), obtained by Wilcoxon rank test of each cluster against all other clusters.

**Supplemental File S2**: Cell type enrichment analysis of cluster classifier genes, showing the inferred identity of clusters of scRNA-seq of haemGx (all conditions). Classifier genes (in Supplemental File S1) were obtained by differential expression in comparison to all other clusters and subjected to enrichment analysis using the PanglaoDB repository (Franzén, Gan and Björkegren, 2019).

**Supplemental File S3**: Summary of projections of scRNA-seq of haemGx cells onto available datasets by Hou et al. (2020), Zhu et al. (2020), and Thambyrajah et al. (2024), showing the identity of each haemGx cell projecting to annotated clusters of the respective studies.

**Supplemental File S4**: List of differentially expressed genes between haemGx-MNX1 and haemGx-MNX1 at 144h and 216h compared to empty vector (EV) control.

### Supplementary Figures

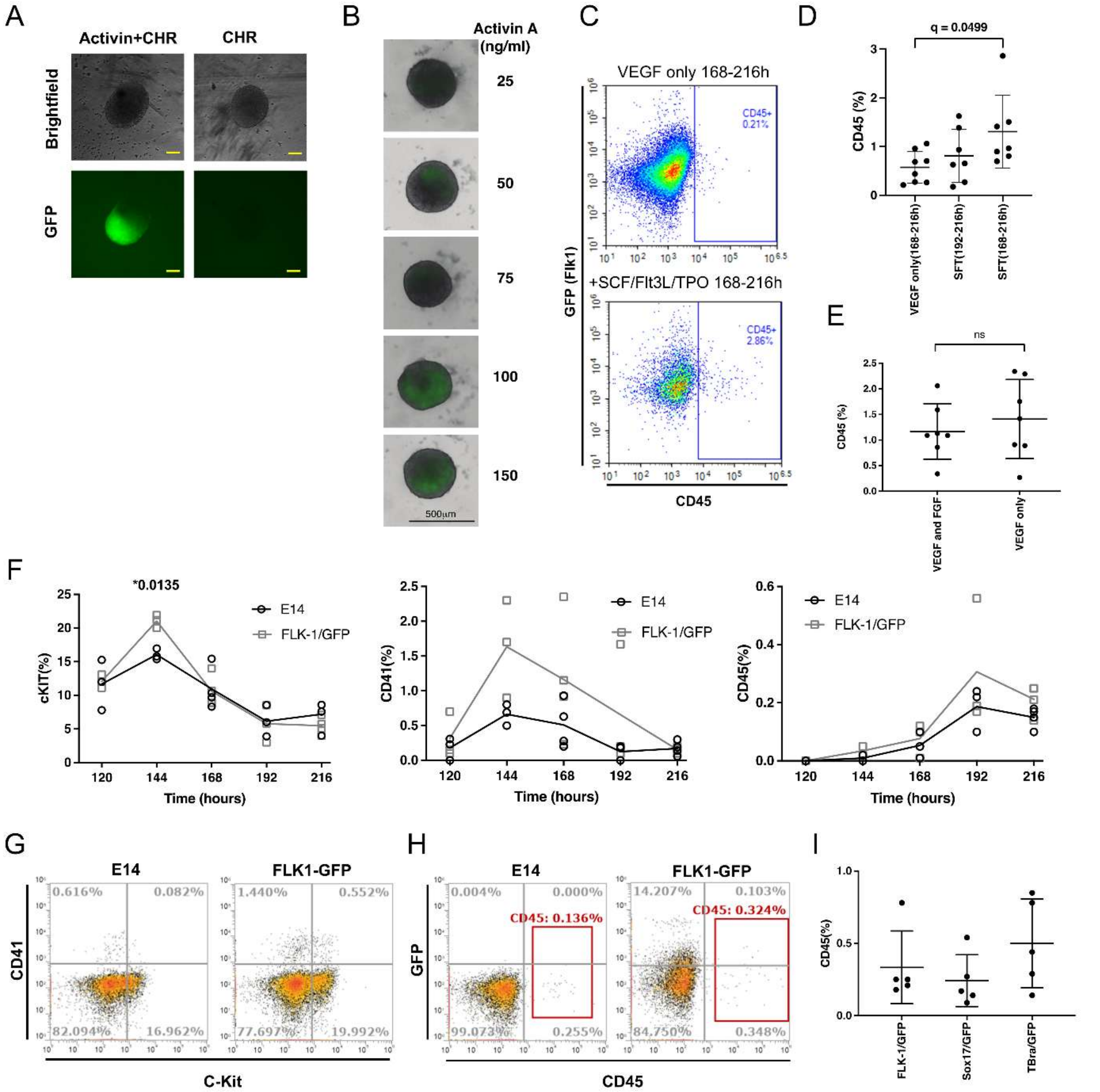

**Figure 1 Supplement 1 – Optimization and reproducibility of the haemGx protocol. (A)** Imaging of a 120h-haemGx with and without the addition of Activin A in conjunction with Chiron 99021 (CHIR) to achieve polarized expression of *Flk-1-GFP*; scale bar: 100mm. **(B)** Titration of Activin A dosage (from 25 ng/ml to 150 ng/ml) required for *Flk-1-GFP* expression activation (green). **(C)** Representative flow cytometry plots of %CD45$^{+}$ cells at 216h without and without addition of SCF, Flt3L and TPO (SFT) from 168h; representative plots. **(D)** Quantification of %CD45$^{+}$ cells by flow cytometry analysis of haemGx cultured with or without SFT, between 192-216h and 168-216h. Mean ± SD of n=7-8 independent experiments; Tukey's multiple

testing with significant q-value<0.05. **(E)** Quantification of %CD45$^+$ cells at 216h by flow cytometry analysis of haemGx cultured in VEGF+FGF or VEGF-only during the 168-216h period. Mean ± SD of n=7 independent experiments; unpaired t-test; ns, not significant. **(F)** Flow cytometry monitoring of C-Kit, CD41, and CD45 markers in haemGx established from E14 and *Flk-1-GFP* mES lines; n=3-4 replicate experiments (n=1, CD45$^+$ 120h). Mixed effects model with Sidak's multiple comparison test; significant adjusted p-value <0.05. **(G)** Representative flow cytometry plots of CD41$^+$ and C-Kit$^+$ cells in E14 and *Flk-1-GFP* mES cell-initiated haemGx. **(H)** Representative flow cytometry plots of CD45$^+$ cells in E14 and *Flk-1-GFP* mES cell-initiated haemGx. **(I)** Quantification of %CD45$^+$ cells in *Flk-1-GFP*, *Sox17-GFP*, and *TBra-GFP* mES cell- initiated haemGx; n=5, mean ± SD. Welch's t-test Sox17-GFP vs. Flk1-GFP, p=0.5254; Tbra-GFP vs. Flk1-GFP, p=0.3782.

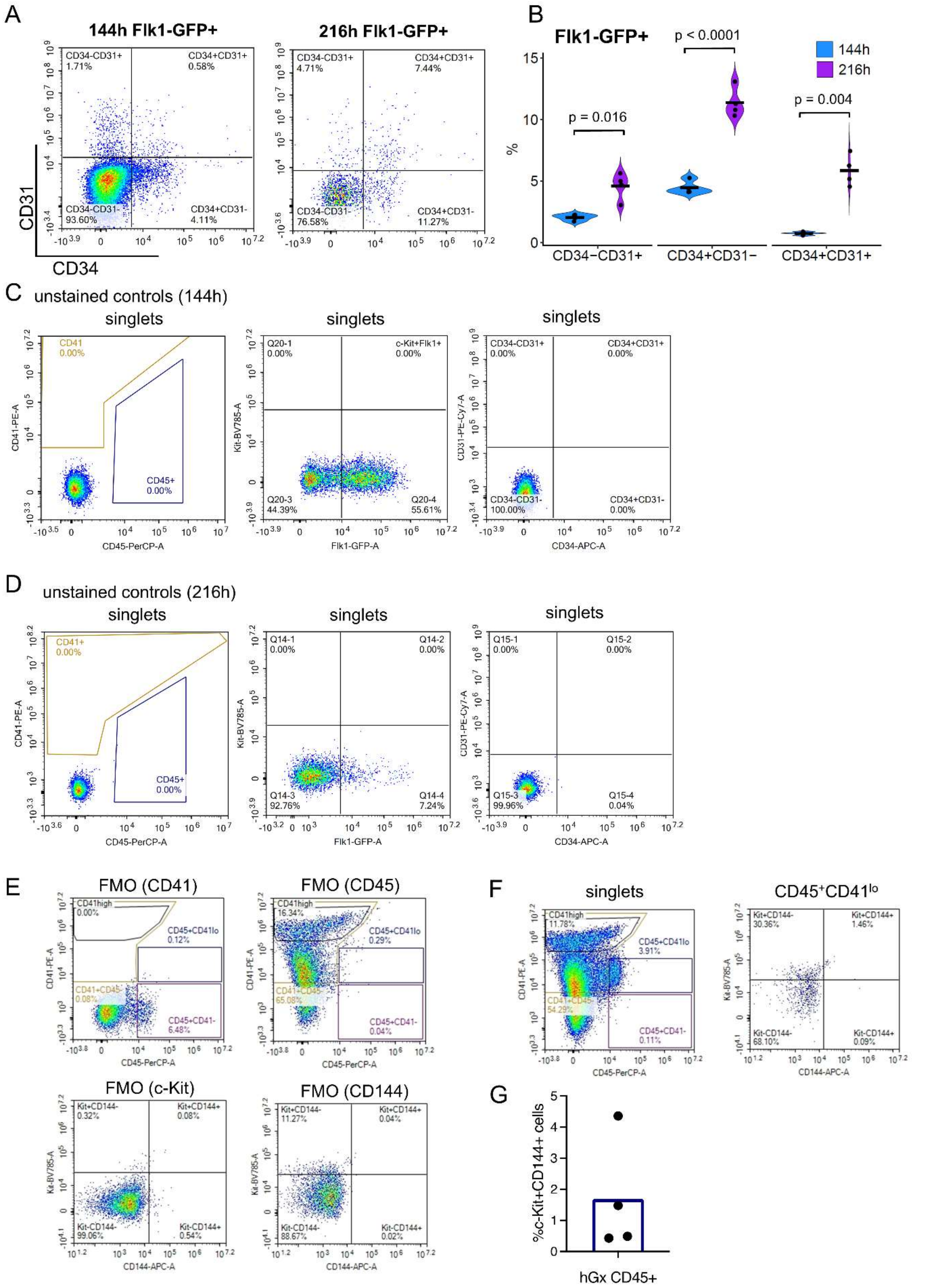


**Figure 2 Supplement 1 – Multi-color flow cytometry detection of surface markers in haemGx. (A)** Representative flow cytometry plots of CD31 and CD34 staining of haemGx at 144h and 216h gated on Flk-1-GFP$^{+}$. **(B)** Quantification of Flk-1-GFP$^{+}$ haemGx populations at

144h and 216h expressing CD34 and CD31. Crossbar shows mean of 4 replicate experiments with individual data points shown; Welch's t-test; p value significant < 0.05. **(C-E)** Unstained controls (C-D) and fluorescence minus one (FMO) controls (E) relative to flow cytometry detection of multi-color of surface markers in Figure 2. **(F)** Representative flow cytometry plot of haemGx of $CD45^{+}$CD41lo fractions co-expressing C-Kit and Cd144 (Ve-Cadherin).

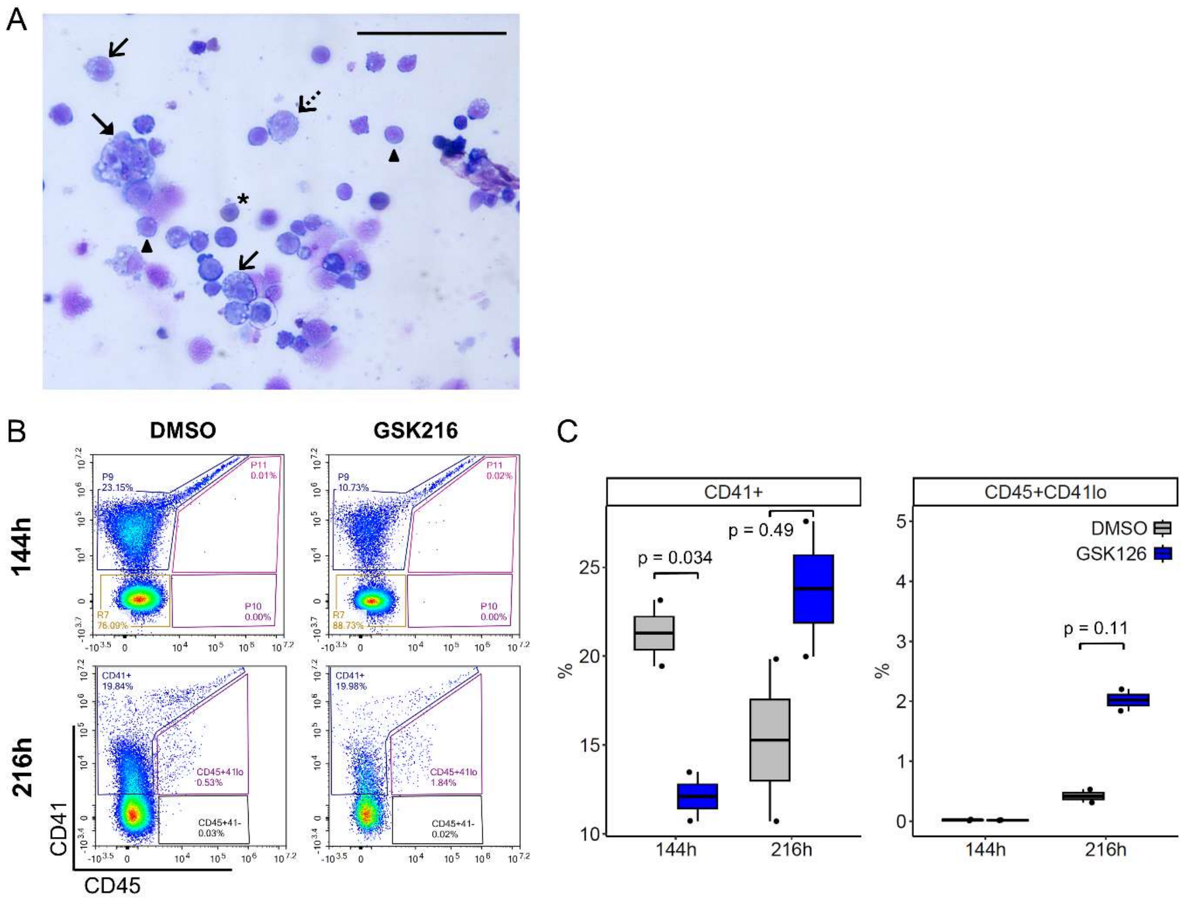


**Figure 2 Supplement 2 – Characterization of hematopoietic output from haemGx. (A)** Representative image of cytospins of dissociated haemGx at 216h stained with Giemsa-Wright's stain. Annotated are cells in the monocytic (dashed open arrow), granulocytic (solid open arrow), megakaryocytic (solid arrow) and erythroid (asterisk) lineages; arrowheads indicate cells with a non-specific blast-like morphology. 40x magnification, scale bar = 100 μm. **(B)** Representative flow cytometry plots of CD41 and CD45 expressing populations in haemGx treated with 0.5 μM EZH2 inhibitor GSK126 or with 0.05% DMSO (control) at 144h and 216h. **(C)** Quantification of flow cytometry analysis in (B) showing the proportion of $CD41^{+}$ and $CD45^{+}CD41^{lo}$ populations in GSK126-treated haemGx. Welch's t-test; p value significant < 0.05.

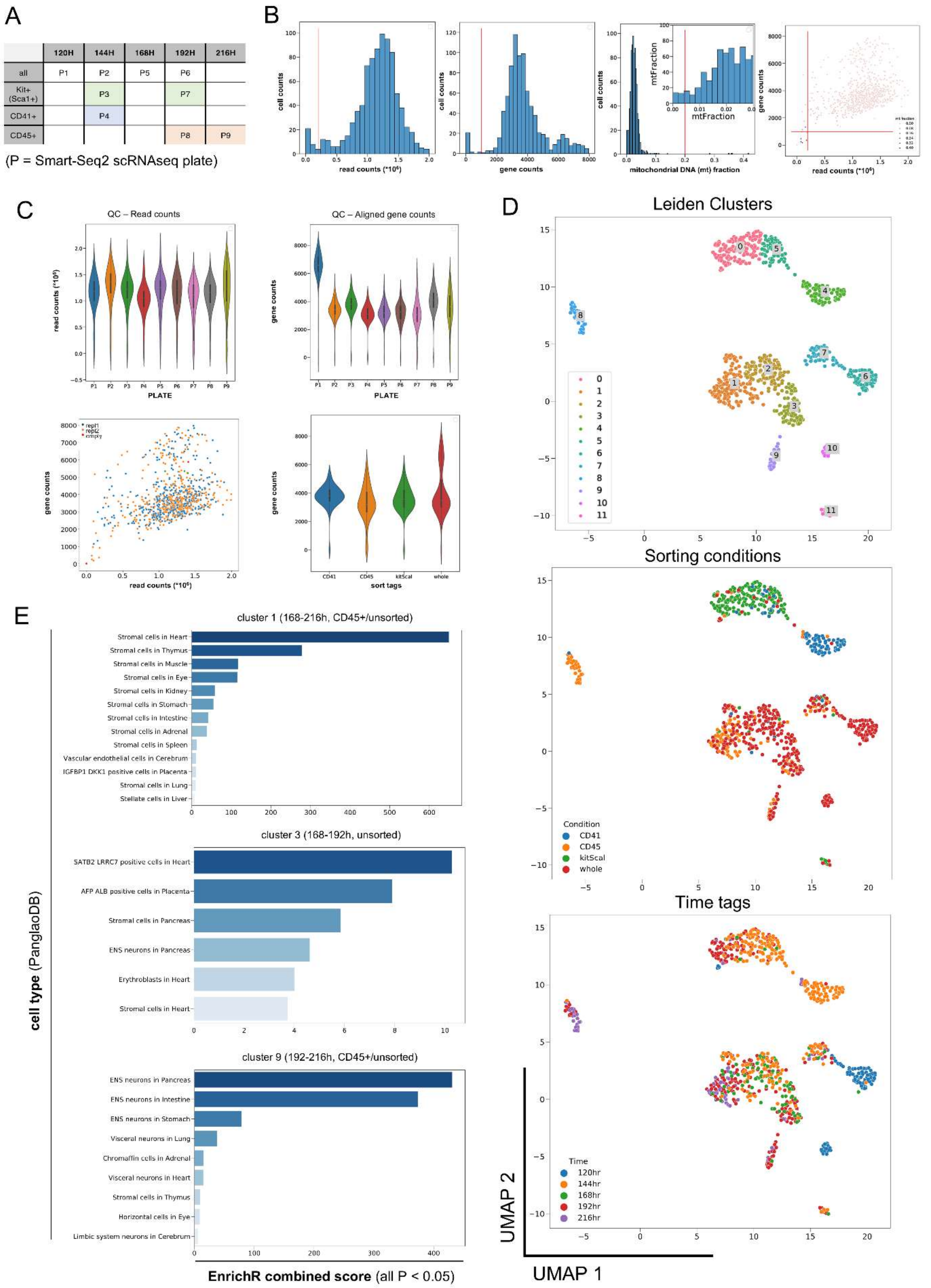


**Figure 3 Supplement 1 – Single-cell-RNAseq (scRNAseq) analysis of haemGx identifies time-dependent signatures of endothelial, hemogenic and stromal cells. (A)** Summary

of plating strategy for scRNA-seq analysis of gastruloids at 120h, 144h, 168h, 192h, and 216h without selection of surface markers (‘all’) or sorted as C-Kit/Sca1+ (green shading), CD41+ (blue), or CD45+ (orange) cells; P = plate. **(B)** ScRNA-seq quality control measures (quantification of mapped reads, individual genes and mapping of reads on to mitochondrial (mt)DNA. **(C)** Mapped read and gene counts for individual Smart-seq2 libraries (plate, P), biological replicate (repl) populations of single cells, and read and map distributions across unsorted and cell surface phenotype-sorted cells. **(D)** UMAP projection of all sequenced cells colored by annotated clusters (left), by sorting markers C-Kit/ScaI, CD41, and CD45, or unfractionated cells (‘whole’ corresponding to ‘all’ in panel A) (center), and by haemGx culture time (right). **(E)** Cell type enrichment analysis of cluster classifier genes, using the PanglaoDB repository (Franzén, Gan and Björkegren, 2019); classifier genes are obtained by differential expression in comparison to all other clusters. Represented are clusters 1, 3 and 9, which are characteristic of timepoints 168-216h, and capture stromal cells putatively relevant for hemogenic support, including autonomic neurons (cluster 9). The statistical power of representation of individual cell types is expressed as the EnrichR combined score with a p-value threshold of $< 0.05$.

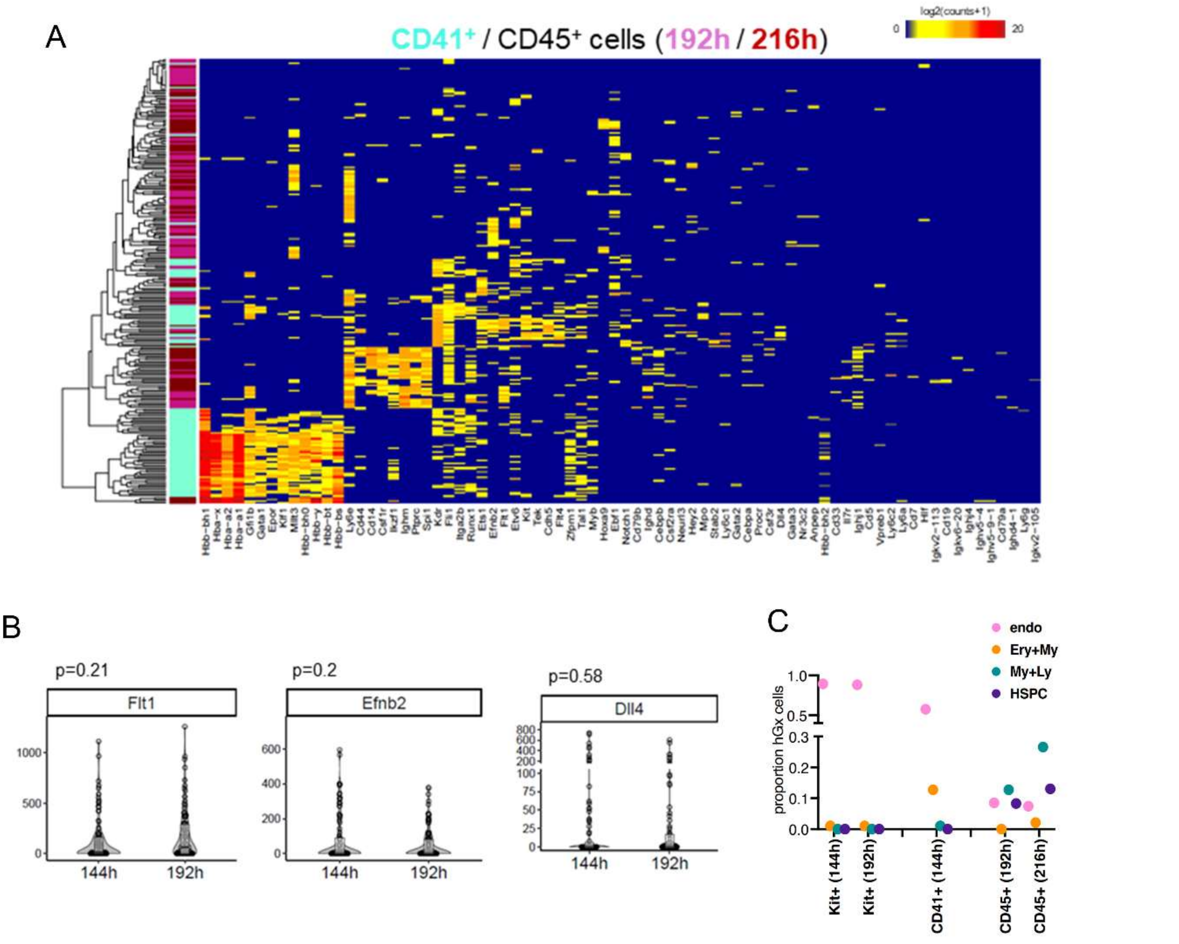


**Figure 3 Supplement 2 – Single-cell-RNAseq (scRNAseq) analysis of haemGx. (A)** Heatmap representation of the expression of endothelial and hematopoietic marker genes in individual haemGx CD41$^{+}$ and CD45$^{+}$ cells sorted at 192 and 216h. Cells and genes ordered by unsupervised hierarchical clustering. **(B)** Violin plots quantifying the expression of arterial markers in C-Kit+ fraction from clusters in Fig. 3A comparing 144h and 192h timepoints. Wilcoxon test; significant p value<0.05. **(C)** Summary representation of the proportion of hGx cells expressing endothelial (endo), erythroid/myeloid (Ery+My), myeloid/lymphoid (My+Ly) and HSPC gene signatures. Endo: *Kdr*; Ery+My: *Epor±Gata1±Klf1±Hbb±Hba±Hbg* + *Spi1±Mpo±Anpep±Csf1/2/3r*); My+Ly: *Spi1±Csf1/2a/3r* + *Ikzf1±Ighm/d±Igk*); HSPC: *Ptprc* + *Myb*).

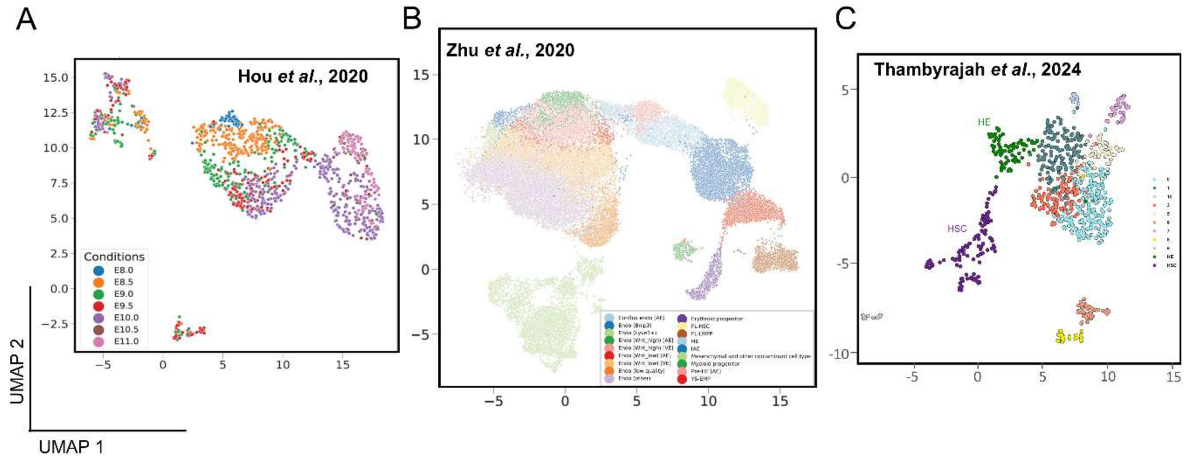


**Figure 4 Supplement 1 – Analysis of mouse embryonic scRNA-seq datasets for transcriptional similarities of haemGx outputs. (A-C)** UMAPs of mouse single-cell RNA-seq datasets annotated as in respective publications: (A) arterial and haemogenic specification in the para-splanchnopleura (pSP) and AGM region between E8.0 and E11 (Hou et al. 2020); (B) YS, AGM and FL progenitors and the AGM EHT (Zhu et al., 2020); (C) HSC emergence from the dorsal aorta HE (Thambyrajah et al., 2024).

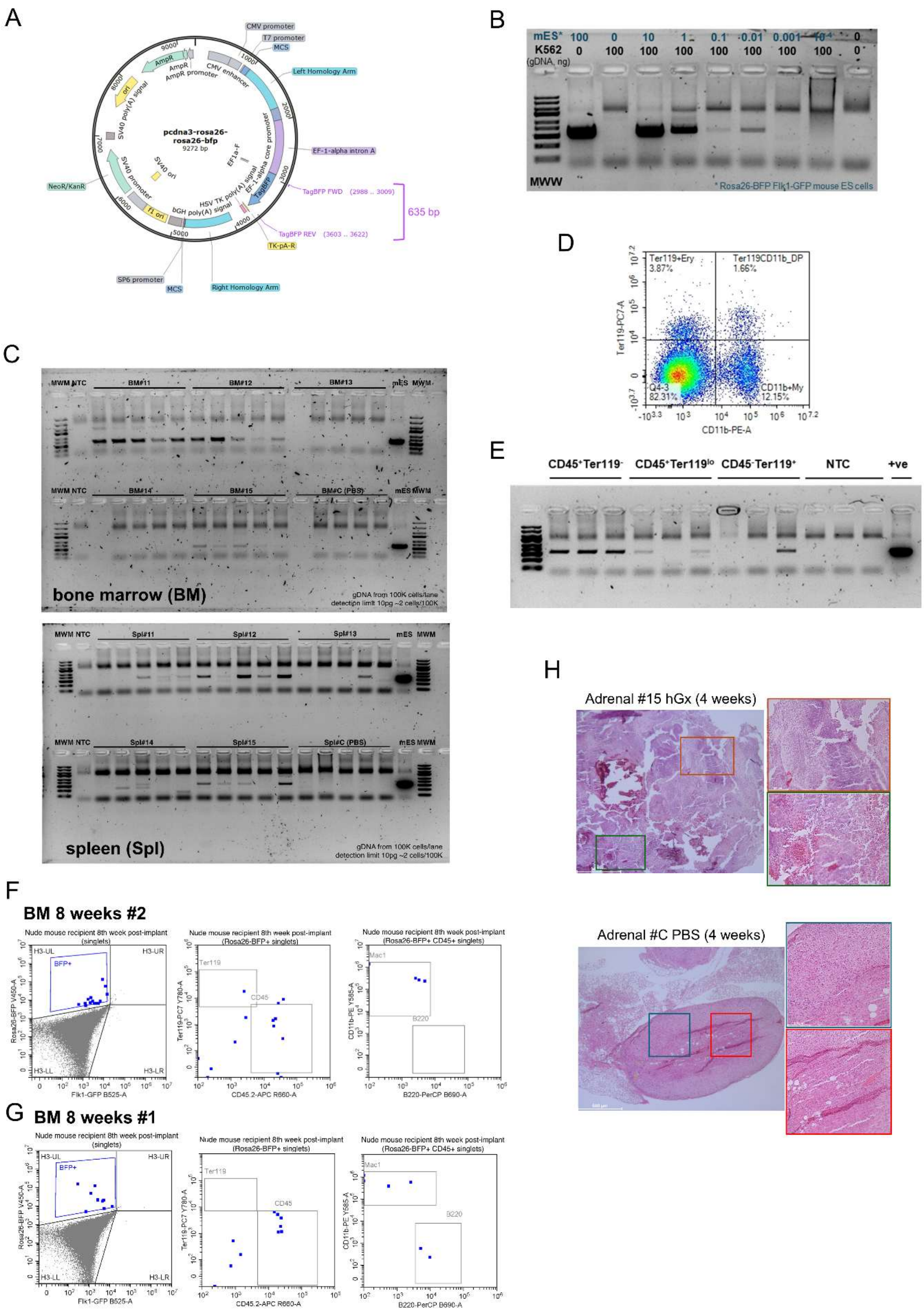
A
pcdna3-rosa26-rosa26-bfp
9272 bp
CMV promoter
T7 promoter
MCS
Left Homology Arm
EF-1-alpha intron A
TagBFP FWD (2988 .. 3009)
635 bp
TagBFP REV (3603 .. 3622)
TK-pA-R
Right Homology Arm
MCS
SP6 promoter
NeoR/KanR
B
mES*
K562
(gDNA, ng)
MWW
* Rosa26-BFP Flk1-GFP mouse ES cells
C
bone marrow (BM)
gDNA from 100K cells/lane
detection limit 10pg ~2 cells/100K
spleen (Spl)
D
Ter119+Ery
3.87%
Ter119CD11b_DP
1.66%
Q4-3
82.31%
CD11b+My
12.15%
Ter119-PC7-A
CD11b-PE-A
E
CD45+Ter119-
CD45+Ter119lo
CD45-Ter119+
NTC
+ve
F
BM 8 weeks #2
G
BM 8 weeks #1
Nude mouse recipient 8th week post-implant
H
Adrenal #15 hGx (4 weeks)
Adrenal #C PBS (4 weeks)

**Figure 4 Supplement 2 – Detection of engraftment potential of late-stage haemGx in adrenal glands of Nude mice. (A)** Map of the *Rosa26-BFP* construct used to target *Flk1-GFP* mES cells employed in implant assays. Primers used for gDNA PCR analysis are indicated. **(B)** PCR detection limit of the BFP amplicon using the primers in (A), in a serial dilution of *Rosa26-BFP*::*Flk1-GFP* mES gDNA into human leukemia cell line K562 gDNA; total 100ng DNA per lane. **(C)** Uncropped image of agarose gel electrophoresis of BFP detection shown in Fig. 4C. **(D)** Representative flow cytometry plot of the spleen of immunodeficient mouse recipients, showing myelo-lymphoid ($CD45^{+}Ter119^{-}$) and erythroid ($Ter119^{+}CD45^{+/-}$) cell fractions at 4 weeks post implantation. **(E)** PCR analysis of *Rosa26-BFP*::*Flk1-GFP* gDNA in the cell fractions in (D) of implanted animal #15 at 4 weeks post-implantation. PCR run in triplicate material from each sample; positive and NTC controls as per Fig. 4C. **(F-G)** Representative flow cytometry interrogation of BFP detection within lineage compartments in the bone marrow (BM) of control and implanted recipient animal #2 (F) and animal #1 (G) at 8 weeks after hGx implantation. **(H)** Histology of implanted and control adrenal glands; paraffin sections stained with hematoxylin & eosin (H&E). Inserts in implanted animal highlight neural (orange) and hematopoietic and renal tubular epithelium (green); inserts in control animal highlight normal cortical (blue) and medullar (red) adrenal architecture, absent in implanted adrenal glands.

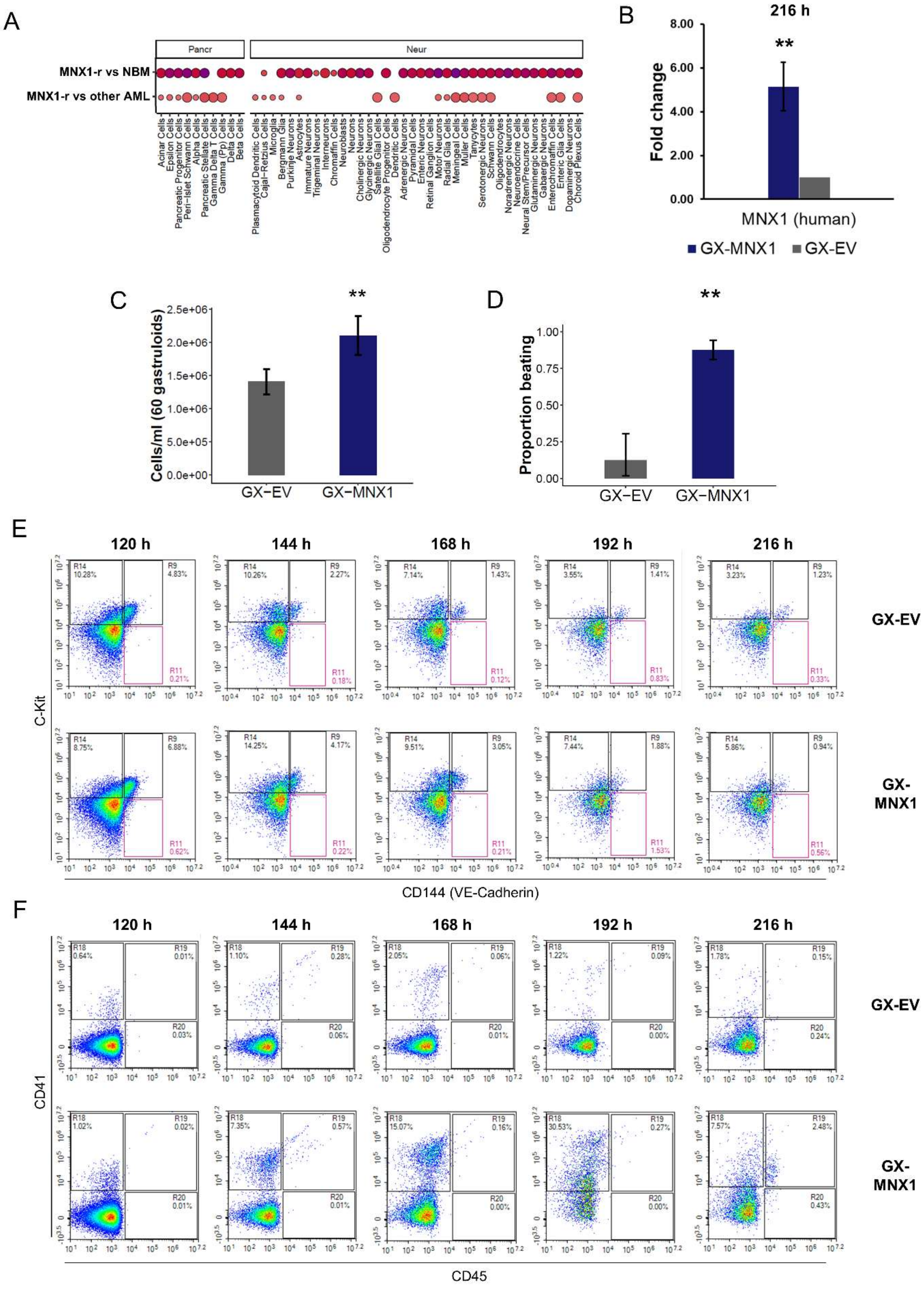
A
Pancr
Neur
MNX1-r vs NBM
MNX1-r vs other AML
Acinar Cells
Epsilon Cells
Pancreatic Progenitor Cells
Peri-islet Schwann Cells
Alpha Cells
Pancreatic Stellate Cells
Gamma Delta T Cells
Gamma (Pp) Cells
Delta Cells
Beta Cells
Plasmacytoid Dendritic Cells
Cajal-Retzius Cells
Microglia
Bergmann Glia
Purkinje Neurons
Astrocytes
Immature Neurons
Trigeminal Neurons
Interneurons
Chromaffin Cells
Neuroblasts
Neurons
Cholinergic Neurons
Glycinergic Neurons
Satellite Glial Cells
Oligodendrocyte Progenitor Cells
Dendritic Cells
Adrenergic Neurons
Pyramidal Cells
Enteric Neurons
Retinal Ganglion Cells
Motor Neurons
Radial Glia Cells
Meningeal Cells
Muller Cells
Tanycytes
Serotonergic Neurons
Schwann Cells
Oligodendrocytes
Noradrenergic Neurons
Neuroendocrine Cells
Neural Stem/Precursor Cells
Glutaminergic Neurons
Gabaergic Neurons
Enterochromaffin Cells
Enteric Glia Cells
Dopaminergic Neurons
Choroid Plexus Cells
B
216 h
**
Fold change
8.00
6.00
4.00
2.00
0.00
MNX1 (human)
GX-MNX1
GX-EV
C
**
Cells/ml (60 gastruloids)
2.5e+06
2.0e+06
1.5e+06
1.0e+06
5.0e+05
0.0e+00
GX−EV
GX−MNX1
D
**
Proportion beating
1.00
0.75
0.50
0.25
0.00
GX−EV
GX−MNX1
E
120 h
144 h
168 h
192 h
216 h
GX-EV
GX-MNX1
C-Kit
CD144 (VE-Cadherin)
F
120 h
144 h
168 h
192 h
216 h
GX-EV
GX-MNX1
CD41
CD45

**Figure 5 Supplement 1 – MNX1 overexpression promotes hemogenic specification in haemGx**. **(A)** Cell type enrichment analysis in MNX1-r AML transcriptomes, compared to normal paediatric bone marrow (BM) or other paediatric AML from the TARGET database. GSEA used representative cell type gene sets from the 2021 DB database; bubble plot shows NES scores (color gradient) and statistical significance (–log10(FDR), bubble size). **(B)** Quantitative (q)RT-PCR analysis of *MNX1* overexpression in 216h-haemGx. Gene expression fold change calculated by normalization to HPRT1. Mean ± SD of 3 replicates; 2-tailed t-test, $p< 0.001$ (**). **(C)** Cell counts of disassembled haemGx at 216h. Mean ± SD of 3 replicate experiments; 2-tailed t-test, $p<0.001$ (**). **(D)** Proportion of EV and MNX1 haemGx exhibiting spontaneous unifocal contractility at 192h, observed in commercial N2B27 medium (see Experimental Procedures); mean ± SD of 3 replicate experiments; 2-tailed t-test, $p<0.001$ (**). **(E-F)** Flow cytometry analysis of specification of **(E)** hemato-endothelial – C-Kit$^+$ and VE-Cadherin$^+$ – and **(F)** hematopoietic progenitor – CD41$^+$ and CD45$^+$ – cells over a 120-216h timecourse of EV and MNX1 haemGx cultures; representative plots.

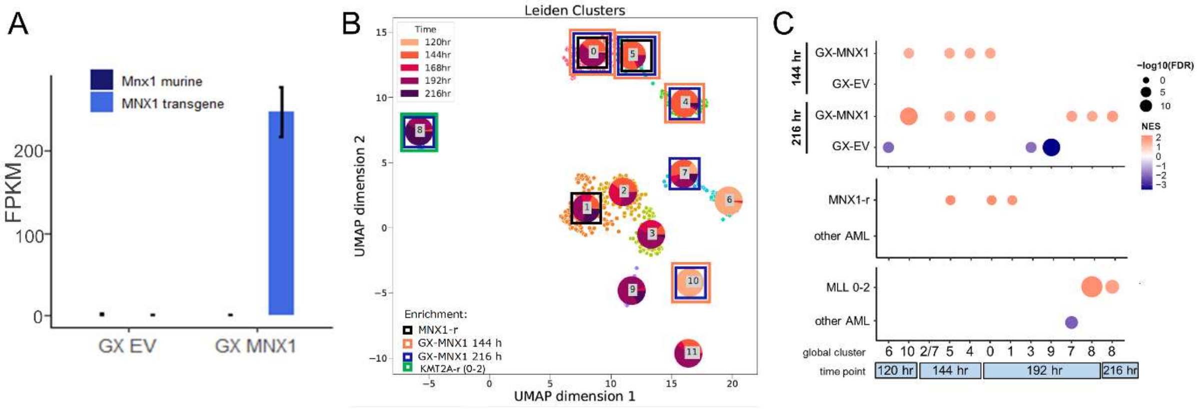


**Figure 5 Supplement 2 – Transcriptional analysis of haemGx with MNX1 overexpression.** **(A)** Expression of human *MNX1* and murine *Mnx1* genes in FPKM units from RNA-seq of MNX1 and EV haemGx. **(B)** UMAP of time-resolved global clustering of scRNA-seq of haemGx showing the enriched clusters in GX-MNX1 at 144 hr (orange boxes) and 216 hr (blue boxes), and MNX1-*r* AML (black boxes) or MLL-AML (green boxes) determined by GSEA. **(C)** Bubble plot of GSEA NES values and statistical significance by –log10(FDR) of enrichments in specific time clusters and corresponding global clusters in GX-MNX1 at 144 hr and 216 hr compared to respective GX-EV, and MNX1-*r* or MLL AML samples compared to other pediatric AML.

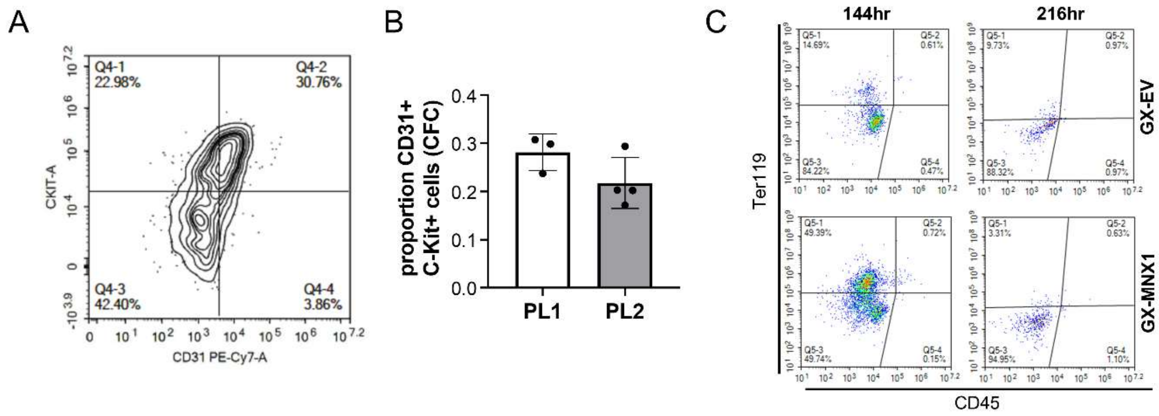


**Figure 6 Supplement 1 – Characterization of MNX1-overexpressing replating cells from haemGx. (A-B)** Representative flow cytometry plot of CD31 and C-Kit staining of CFC plating of GX-MNX1 (left) and quantification of sustained expression of $CD31^{+}CKit^{+}$ cells through replatings. **(C)** Representative flow cytometry plots of serially re-plating MNX1 cells from 144h and 216h-gastruloids (plate 5 cells) stained with Ter119 and CD45.

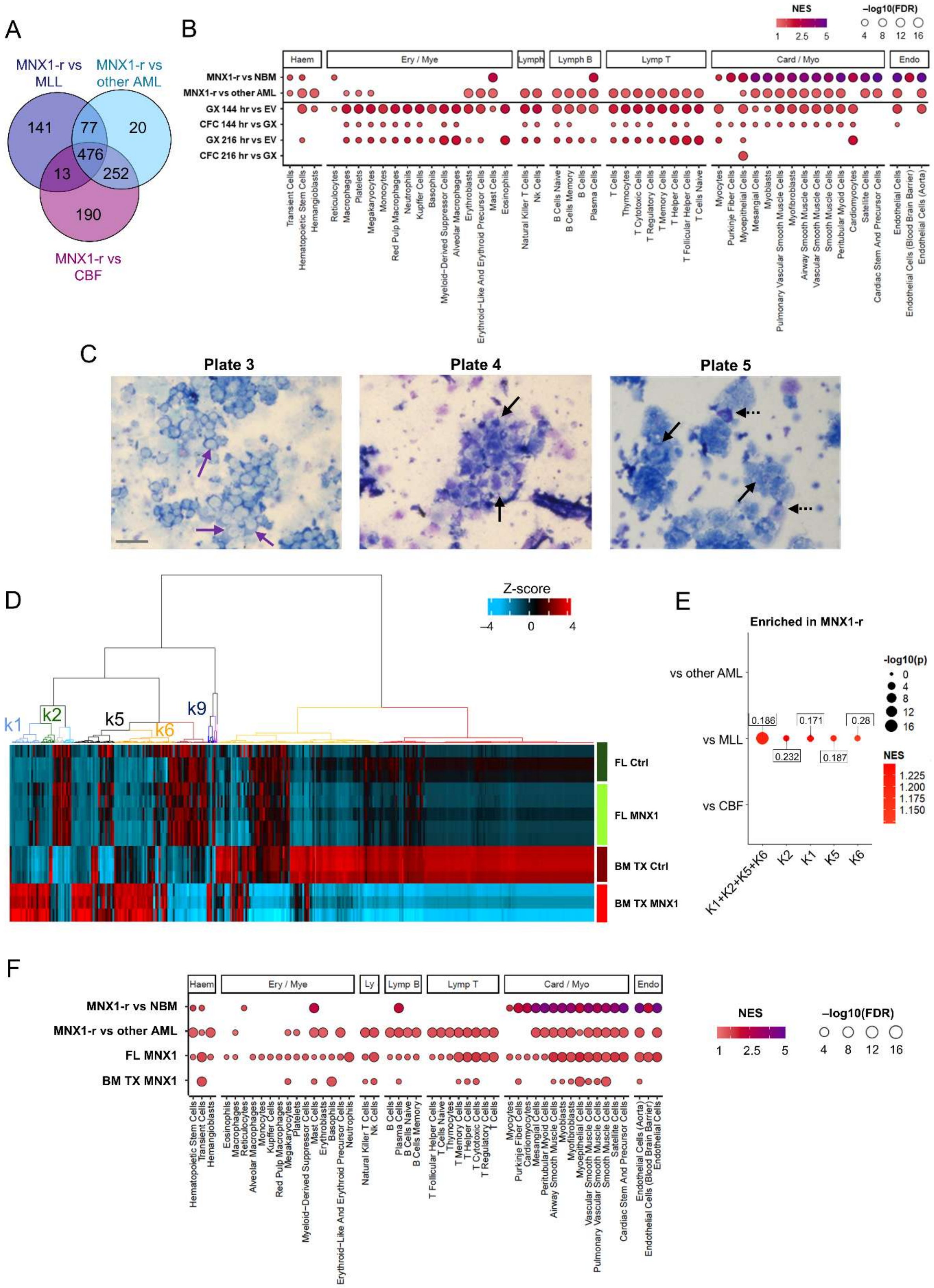
A
MNX1-r vs MLL
MNX1-r vs other AML
MNX1-r vs CBF
141
77
20
476
13
252
190
B
NES
1
2.5
5
–log10(FDR)
4
8
12
16
Haem
Ery / Mye
Lymph
Lymph B
Lymp T
Card / Myo
Endo
MNX1-r vs NBM
MNX1-r vs other AML
GX 144 hr vs EV
CFC 144 hr vs GX
GX 216 hr vs EV
CFC 216 hr vs GX
Transient Cells
Hematopoietic Stem Cells
Hemangioblasts
Reticulocytes
Macrophages
Platelets
Megakaryocytes
Monocytes
Red Pulp Macrophages
Neutrophils
Kupffer Cells
Basophils
Myeloid−Derived Suppressor Cells
Alveolar Macrophages
Erythroblasts
Erythroid−Like And Erythroid Precursor Cells
Mast Cells
Eosinophils
Natural Killer T Cells
Nk Cells
B Cells Naive
B Cells Memory
B Cells
Plasma Cells
T Cells
Thymocytes
T Cytotoxic Cells
T Regulatory Cells
T Memory Cells
T Helper Cells
T Follicular Helper Cells
T Cells Naive
Myocytes
Purkinje Fiber Cells
Myoepithelial Cells
Mesangial Cells
Myoblasts
Pulmonary Vascular Smooth Muscle Cells
Myofibroblasts
Airway Smooth Muscle Cells
Vascular Smooth Muscle Cells
Smooth Muscle Cells
Peritubular Myoid Cells
Cardiomyocytes
Satellite Cells
Cardiac Stem And Precursor Cells
Endothelial Cells
Endothelial Cells (Blood Brain Barrier)
Endothelial Cells (Aorta)
C
Plate 3
Plate 4
Plate 5
D
Z-score
–4
0
4
k1
k2
k5
k6
k9
FL Ctrl
FL MNX1
BM TX Ctrl
BM TX MNX1
E
Enriched in MNX1-r
vs other AML
vs MLL
vs CBF
0.186
0.232
0.171
0.187
0.28
-log10(p)
0
4
8
12
16
NES
1.225
1.200
1.175
1.150
K1+K2+K5+K6
K2
K1
K5
K6
F
Haem
Ery / Mye
Ly
Lymp B
Lymp T
Card / Myo
Endo
MNX1-r vs NBM
MNX1-r vs other AML
FL MNX1
BM TX MNX1
NES
1
2.5
5
–log10(FDR)
4
8
12
16
Hematopoietic Stem Cells
Transient Cells
Hemangioblasts
Eosinophils
Macrophages
Reticulocytes
Alveolar Macrophages
Monocytes
Kupffer Cells
Red Pulp Macrophages
Megakaryocytes
Platelets
Myeloid−Derived Suppressor Cells
Mast Cells
Erythroblasts
Basophils
Erythroid−Like And Erythroid Precursor Cells
Neutrophils
Natural Killer T Cells
Nk Cells
B Cells
Plasma Cells
B Cells Naive
B Cells Memory
T Follicular Helper Cells
T Cells Naive
Thymocytes
T Memory Cells
T Helper Cells
T Cytotoxic Cells
T Regulatory Cells
T Cells
Myocytes
Purkinje Fiber Cells
Cardiomyocytes
Mesangial Cells
Peritubular Myoid Cells
Airway Smooth Muscle Cells
Myoblasts
Myofibroblasts
Myoepithelial Cells
Vascular Smooth Muscle Cells
Pulmonary Vascular Smooth Muscle Cells
Smooth Muscle Cells
Satellite Cells
Cardiac Stem And Precursor Cells
Endothelial Cells (Aorta)
Endothelial Cells (Blood Brain Barrier)
Endothelial Cells

**Figure 7 Supplement 1 – Transcriptional analyses of haemGx with *MNX1* overexpression compared to patient signatures and current *in vivo* model. (A)** Venn diagram showing the intersection of leading-edge genes enriched in *MNX1*-r patient vs MLL, core-binding factors (CBF), or other pediatric AML from GSEA analysis using the K14 signature. **(B)** Cell type enrichment analysis in *MNX1*-r AML transcriptomes, compared to normal paediatric bone marrow (BM) or other paediatric AML from the TARGET database, with comparison with transcriptomes of haemGx and CFCs at from MNX1 vs EV conditions at 144h and 216h. GSEA used representative cell type gene sets from the Panglao DB 2021database; bubble plot shows NES scores (color gradient) and statistical significance (–log10(FDR), bubble size). **(C)** Representative Giemsa-Wright stained dissociated CFC replating MNX1 haemGx cells from 144h; purple arrows, undifferentiated blasts; solid black arrow, mast cell precursors; dashed black arrow, mast cells. **(D)** Transcriptional re-analysis of Waraky et al (2024) in vivo model of MNX1-OE leukemia from transplanted fetal liver (FL) cells. Heatmap comparing the expression levels of all DEG between all conditions, i.e. FL control (FL Ctrl) vs FL with MNX1-OE (FL MNX1) in liquid culture, and bone marrow (BM) from mice transplanted with FL Ctrl (BM TX Ctrl) vs leukemic animals transplanted with FL MNX1 cells (BM TX MNX1). Hierarchical clustering by Ward D method on Euclidean distances identifies 12 clusters (k). Clusters k1, k2, k5, k6 and k9 are specifically up-regulated in BM TX MNX1 and putatively represent MNX1-OE leukemia programs. Analysis matches Fig. 7B for MNX1-OE haemGx. **(E)** GSEA plots of significant results (by p value $< 0.05$) for gene set extracted from k1, k2, k5, k6 and k9, individually or combined, from (D) against MNX1-r patients RNA-seq counts vs MLL, core-binding factors (CBF), or other pediatric AML. FDR values are indicated for each enrichment point. Analysis matches Fig. 7C for MNX1-OE haemGx. **(F)** Cell type enrichment analysis in MNX1-r AML transcriptomes, compared to normal pediatric bone marrow (BM) or other pediatric AML from the TARGET database. GSEA used representative cell type gene sets from the Panglao DB 2021 database; bubble plot shows NES scores (color gradient) and statistical significance (–log10(FDR), bubble size). Analysis matches Fig. S6E for MNX1-OE haemGx.
